# The role of phenotypic plasticity in the establishment of range margins

**DOI:** 10.1101/2021.10.04.463099

**Authors:** Martin Eriksson, Marina Rafajlović

## Abstract

It has been argued that adaptive phenotypic plasticity may facilitate range expansions over spatially and temporally variable environments. However, plasticity may induce fitness costs. This may hinder the evolution of plasticity. Earlier modelling studies examined the role of plasticity during range expansions of populations with fixed genetic variance. However, genetic variance evolves in natural populations. This may critically alter model outcomes. We ask: How does the capacity for plasticity in populations with evolving genetic variance alter range margins that populations without the capacity for plasticity are expected to attain? We answered this question using computer simulations and analytical approximations. We found a critical plasticity cost above which the capacity for plasticity has no impact on the expected range of the population. Below the critical cost, by contrast, plasticity facilitates range expansion, extending the range in comparison to that expected for populations without plasticity. We further found that populations may evolve plasticity to buffer temporal environmental fluctuations, but only when the plasticity cost is below the critical cost. Thus, the cost of plasticity is a key factor involved in range expansions of populations with the potential to express plastic response in the adaptive trait.

## 1 Introduction

Due to ongoing climate change and increasing human impact on ecosystems, many populations need to adapt to novel conditions either in their present geographical distributions, or in new areas they face with while altering their ranges [1, 2, 3, 4, 5]. A critical factor constraining local adaptation and thereby precluding successful range expansions is maladaptive gene flow [6, 7]. Theoretically, it has been shown that, when genetic variance is fixed and the population is faced with a sufficiently steep constant environmental gradient, maladaptive gene flow swamps local adaptation This results in a finite range of the population [8] (see also [9]).

However, genetic variance in natural populations is expected to evolve. Notably, the above theoretical prediction is critically altered when genetic variance is allowed to evolve. Under this assumption, populations expanding their ranges over an environment that changes linearly in space (with a constant carrying capacity) will either adapt to the entire available habitat or face global extinction [10]. In this case, thus, range margins are trivial: they either coincide with the habitat edges or, when the habitat is unlimited, range margins are absent.

By contrast, non-trivial range margins exist when a population expands its range over a steepening environmental gradient, and this is true even when the available habitat is infinite [10, 11]. In this case, local genetic variance increases with increasing local steepness of the environmental gradient until the genetic load becomes so strong that the population is precluded from adapting further. This is seen as a progressively decreasing expected local population size (despite the assumption that the carrying capacity is constant over the habitat) down to the point where drift becomes stronger than selection [11]. Conversely, for range expansions over environments that change linearly in space (with genetic variance allowed to evolve), drift may cause non-trivial range margins to be established when the local carrying capacity decreases away from the core habitat [11].

The results outlined above deliver an insight into potential mechanisms involved in the establishment of range limits. However, they do not account for phenotypic plasticity (hereafter referred to as *plasticity*), that is, the ability of a genotype to produce different phenotypes depending on the environment [12, 13, 14, 15, 16].

Plasticity may be an important mechanism for populations to buffer environmental changes, as shown both empirically [17, 18, 19, 20, 21, 22] and theoretically [9, 13, 16, 23, 24, 25]. This is especially true when plasticity is adaptive (moving phenotypes towards the local optimum) [26, 27]. However, plasticity may also be neutral or maladaptive (moving phenotypes away from the local optimum) [28]. Maladaptive plasticity may have a temporary adverse effect on local adaptation but, in the long run, it may promote genetic adaptation by enhancing the strength of selection [29, 30, 31].

However, it has been empirically observed that plasticity does not always contribute to the persistence of populations [32]. Indeed, plasticity may have costs or limits [33, 34], and these may limit the utility of plasticity for adaptation to new or changing environments [35].

Understanding the evolution of plasticity along environmental gradients, and its role on local adaptation has been the focus of a number of theoretical studies (e.g., [9, 23, 25, 36]). For example, in [23], it was found that, in areas where the difference between the local phenotypic optimum and the globally average optimum was larger, local adaptation was facilitated by the evolution of locally higher plasticity. This is, in part, because migration was implemented according to the island model (*sensu* [37]). In this model, immigration has a strongly deleterious effect on the local mean phenotype when it deviates strongly from the global mean. This causes local maladatation, which produces directional selection to restore the local mean phenotype to its optimum. Consequently, plasticity is under stronger selection when the difference between the local environment and the reference environment (as defined in [38]) is larger. Notably, the model in [23] was deterministic and it was assumed that genetic variance was fixed. These assumptions may bear both qualitative and quantitative consequences on the results obtained.

A similar result was found in a model with an environment that changes linearly in space and a density regulated population (*albeit* without drift) [9]. As a consequence, plasticity increased the range attained by the population in comparison to the case without plasticity [9]. Notably, the results in [9] relied on two assumptions that may critically affect the model outcomes, especially regarding the range that the population is expected to attain. Namely, genetic variance was fixed and the carrying capacity was decreasing away from the centre of the range. As explained above (see also [11]), these assumptions are responsible for the establishment of non-trivial range margins in an environment that changes linearly in space. These assumptions were relaxed in [25], where it was found that transiently increased plasticity evolves in spatial locations that have a long history of environmental change, or at the expansion front for a population undergoing range expansion into a habitat that requires new adaptations (termed “niche expansion” in that study). Notably, in [25] the environment changed linearly in space. This precluded the establishment of non-trivial range margins in that study.

In summary, the role of plasticity on the establishment of non-trivial range margins, when genetic variance is allowed to evolve, remains unclear. Here we address this issue by modelling a population, with evolving genetic variance, expanding its range over a steepening environmental gradient. This is a situation in which a population without plasticity is expected to attain a non-trivial range margin, even when the carrying capacity is not constrained to be decreasing away from the core habitat [11]. Specifically, we ask: How does a population’s capacity for plasticity impact on the establishment of range margins when genetic variance is allowed to evolve and the local carrying capacity is constant? What is the role of plasticity costs in this context? What is the spatial pattern of allele frequencies at the underlying loci?

To answer these questions, we extend the individual-based model from [11] to encompass the capacity for plasticity. This was done by assuming that the adaptive trait had a non-plastic and a plastic component. We further used a simplified version of our model to derive an analytical expression for the *optimal plasticity*, that is plasticity that maximises the population’s mean fitness in quasi-equilibrium. We note that we used here *quasi*, because all finite populations with a finite growth rate will eventually go extinct [39]. With this caution in mind, we use throughout *equilibrium* in place of *quasi-equilibrium*, for simplicity.

Our main finding is that there is a critical cost of plasticity below which the ability to express and evolve plasticity leads to a wider range than for populations lacking this ability. Furthermore, we found a second critical cost below which the range may be infinite. Finally, we found that the equilibrium spatial patterns of allele frequencies at loci contributing to the non-plastic component of the phenotype have the same clinal shape as without plasticity, but the spacing between the clines is increased when plasticity is larger. For the plastic component of the phenotype, we found that the frequencies of alleles associated with positive plasticity increased in a cline-like manner towards the edges of the habitat only when the cost of plasticity was below the critical cost. Otherwise no clinal pattern emerged.

## 2 Methods

We used computer simulations to investigate the impact of plasticity on the evolution of range margins. The simulations were performed using custom-made Matlab code (will be submitted to Dryad upon acceptance of the manuscript).

We extended the model previously considered in [40] (see also [11, 41]), in which a population expanded its range over a habitat with a steepening environmental gradient, assuming a single trait under selection. In addition, in the present work we assumed that the phenotype was determined by a combination of a non-plastic and a plastic component. We further allowed the optimal phenotype to fluctuate in time. These model modifications are explained in more detail below.

The habitat consisted of a one-dimensional chain of *M* = 220 demes, each with a local carrying capacity of *K* = 100 diploid individuals (unless otherwise stated; see Appendix A for details regarding parameter choices, and table 1 that lists the notations used throughout). The generations were discrete and non-overlapping. The individuals were monoecious and mating was assumed to occur randomly with selfing allowed at no cost. As in [11, 40], we assumed a gradually steepening environmental gradient along the habitat: in each deme, *i* = 1, 2, …, *M*, the average optimal phenotype for the trait under selection, 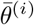, was given by a cubic polynomial of the deme number, *i*, such that 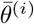 ranged between ±252.9 (figure A1). This polynomial was chosen to be symmetric with a horizontal inflection point at the centre of the habitat, where the optimal phenotype was assumed to be zero (Appendix A). Recall that a steepening (but not a constant) environmental gradient allows non-trivial range margins to be established in a population lacking the capacity for a plastic response. To further understand the role of a gradually steepening as opposed to a constant gradient on the evolution of the spatial pattern in plasticity in the population, we also performed simulations along an environment that changes linearly in space (i.e., along a constant gradient; Appendix A). We further assumed that the realised optimal value for the phenotype is either temporally constant or that it fluctuates in time. In the latter case, we assumed that in deme *i* in generation *τ*, the optimal phenotype (denoted by 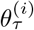 hereafter) is a normally distributed random variable with mean 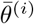 and standard deviation *σ*_*θ*_ (see table A1 for a list of parameter values explored). For simplicity, we assumed that fluctuations in the optimal phenotype were temporally and spatially uncorrelated.

**Table 1:** Explanation of the notations used throughout.

| Notation | Description |
| --- | --- |
| $M$ | Number of demes in the habitat |
| $K$ | Carrying capacity per deme |
| $N_\tau^{(i)}$ | Local population size in deme $i$ in generation $\tau$ |
| $\theta_\tau^{(i)}$ | Optimal phenotype in deme $i$ in generation $\tau$ |
| $\bar{\theta}^{(i)}$ | Average optimal phenotype in deme $i$ |
| $\sigma_\theta$ | Standard deviation of environmental fluctuations |
| $u_{\tau,k}^{(i)}$ | Phenotype of the trait under selection for individual $k$<br>in deme $i$ in generation $\tau$ |
| $z_{\tau,k}^{(i)}$ | Non-plastic component of the phenotype for<br>individual $k$ in deme $i$ in generation $\tau$ |
| $g_{\tau,k}^{(i)}$ | Plasticity of the phenotype for individual $k$<br>in deme $i$ in generation $\tau$ |
| $W_{\tau,k}^{(i)}$ | Fitness of individual $k$ in deme $i$ in generation $\tau$ |
| $C_\gamma(g_{\tau,k}^{(i)}, \delta)$ | Cost-related function for plasticity |
| $\gamma$ | Shape parameter for the cost-related function |
| $\delta$ | Scale parameter for the cost-related function |
| $r_{\tau,k}^{(i)}$ | Growth rate of individual $k$ in deme $i$ in generation $\tau$ |
| $r_m$ | Maximal intrinsic growth rate |
| $V_S$ | Width of stabilising selection |
| $\mu$ | Mutation rate |
| $L$ | Number of loci under selection for the non-plastic as well as<br>for the plastic component of the phenotype (total of $2L$ loci) |
| $\alpha$ | Effect size of alleles coding for the non-plastic component<br>of the phenotype |
| $\beta$ | Effect size of alleles coding for plasticity |
| $s$ | Selection per locus for loci underlying the non-plastic<br>component of the phenotype, $s = \alpha^2/(2V_S)$ |
| $\sigma$ | Standard deviation of Gaussian dispersal function |

We assumed that the phenotype, 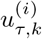, of the trait under selection for individual *k* in deme *i* in generation *τ* was equal to the sum of a non-plastic and a plastic component

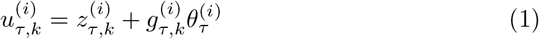

where 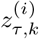 denotes the non-plastic component and 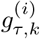 denotes the magnitude of the individual’s plastic response relative to the local phenotypic optimum (hereafter referred to as *plasticity*). The full plastic component of the phenotype was assumed to be equal to 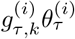, reflecting a common assumption (e.g., [24, 25]) that the same environmental variable determines both the plastic response and the optimal phenotype. For simplicity, we use 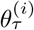 to denote both the optimal phenotype and the environmental cue that affects the plastic response. Note that 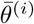 was zero in the centre of the habitat, hence plasticity had, on average (i.e., ignoring the temporal fluctuations), no effect on the average phenotype there. This setting corresponds to treating the centre of the habitat (which is the source of expansion in the model) as the *reference environment* for the plastic response [38]. Note that equation (1) corresponds to equation (2) in [25] in the special case when the reference environment, *g*_2_, in the notation from [25], is zero.

In our model, the non-plastic component of the phenotype, 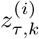, and plasticity 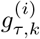, were each underlain by *L* freely recombining bi-allelic loci with additive allele effects, that is, in total there were 2*L* loci under selection (but we also performed simulations where the number of loci for the plastic and non-plastic component were different; Appendix C). The two possible allele effect sizes for the loci underlying 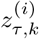 were ±*α/*2 with 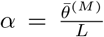 so that in the absence of plasticity (i.e., when 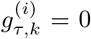), the *L* loci underlying 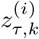 were just enough to constitute the average minimal and maximal optimal phenotypes in the habitat, i.e., the optima at the habitat edges (analogously to [40]). The two possible allele effect sizes for the loci underlying 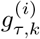 were ±*β/*2 with *β* = 2*/L* so that 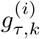 was between −2 and 2. In a special case when 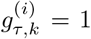 and 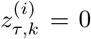, it follows that 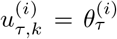. Noting that the optimal phenotype in the source of the expansion is, on average, zero, we refer to plasticity of one (i.e., 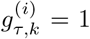) as *perfect plasticity*, because it allows perfect adaptation everywhere without any evolution of the non-plastic component with respect to the source of the expansion.

Apart from assuming that plasticity had a polygenic basis, we also allowed it to be potentially costly. Namely, we modelled the fitness 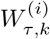 of individual *k* in deme *i* in generation *τ* as

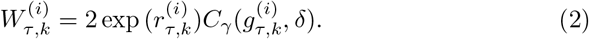

In equation (2), the factor 2 is included due to diploidy, 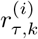 is the growth rate and 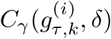 is a cost-related function accounting for a maintenance cost of plasticity (*sensu* [33]), such that costs are larger when 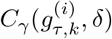 is smaller, and *vice versa*. These components are further explained next.

The growth rate, 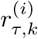, was assumed to be given by

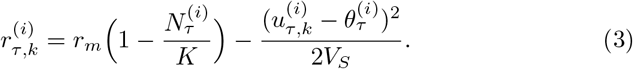

Here, *V*_*S*_ denotes the width of stabilizing selection and we assumed throughout that *V*_*S*_ = 2. Furthermore, *r*_*m*_ denotes the maximal intrinsic growth rate and it was set to *r*_*m*_ = 1 in our simulations. Finally, 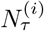 denotes the population size in deme *i* in generation *τ*, and 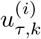 denotes the phenotype, given by equation (1). Note that when 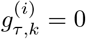 and *σ*_*θ*_ = 0, the model reduces to the one considered in [40]. Our model did not contain any residual component of phenotypic variance caused by environmental factors in addition to the variability in 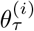.

We assumed that 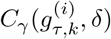 is a decreasing function of the absolute value of plasticity 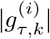 (similarly as in [25]), that is:

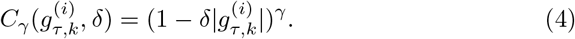

In equation (4), *δ* and *γ* are non-negative parameters, assumed to be constant over time and the same for all individuals. The parameter *δ* determines the threshold plasticity above which the maximal fitness of an individual is non-positive. When 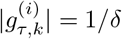, it follows that 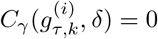, and hence 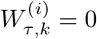. To avoid occurrences of negative fitness, we define 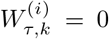 when 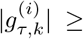 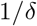. Conversely, the parameter *γ* is a shape parameter, determining whether plasticity costs are more sensitive to high or low plasticity. When *δ* = 0 and/or *γ* = 0, it follows that 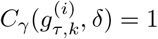, and thus there is no cost of plasticity. The cost of plasticity increases with increasing *δ* and/or *γ* (keeping 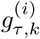 constant). A graphical illustration of the cost-related function 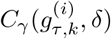 for *γ* = 1 and *γ* = 0.5 is shown in figure A2 in Appendix A.

The life cycle of individuals was modelled as follows. First, each individual contributed a random number of gametes sampled from a Poisson distribution with mean 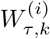 (equation (2)). Plasticity was expressed during the adult life stage in the same environment where the individuals mated. Recombination occurred independently for each gamete, with free recombination between all loci. Second, at each locus mutation occurred reversibly and symmetrically between the two possible alleles with probability *μ* = 10^−6^ per allele, per gamete, per generation. Third, pairs of gametes were chosen uniformly at random to form zygotes (thus, selfing was possible). Finally, the parents were removed and the zygotes dispersed according to a Gaussian function with mean 0 and standard deviation *σ* = 1, as described in [40]. After migration, zygotes were treated as adults.

At the start of each simulation, a fraction of the habitat was occupied, and we initialised genotypes in such a way that the average phenotype of the population followed the local optimum in the occupied demes and all individuals initially had plasticity of zero (Appendix A). Consequently, the (narrow-sense) heritability [42] of the phenotype was initially set to 1 because the total phenotypic variance was governed entirely by the genetic variance of the non-plastic component. However, during the course of simulations, the heritability evolved and potentially varied throughout the range (when plasticity varied spatially), attaining small values when plasticity was large.

After initialising the starting genotypes, we simulated a burn-in period of 100, 000 generations in the source population before we allowed expansion over the empty demes. The burn-in period allowed us to initiate range expansion from an old source population. During the burn-in period the source population stabilised under migration, selection, mutations, drift and possible interactions between the plastic and non-plastic component of the phenotype. This reduced the impact of our choice regarding the starting genotypes (described in Appendix A) on the follow-up dynamics of range expansion.

During the burn-in period, the population was restricted to *M/*5 demes in the centre of the habitat. The boundaries were reflecting, that is, individuals remained at boundary demes instead of dispersing out of the initial range. Note that the number of migrants reaching the boundaries was finite in every generation because all demes have a finite number of individuals prior to migration, and dispersal distance is relatively small (*σ* = 1).

After the burn-in, the population was allowed to expand its range for additional 100, 000 generations (or 200, 000 generations in some cases; Appendix A). As during the burn-in period, the habitat had reflecting boundaries.

We examined different parameter sets, chosen to below, close to, or above the critical cost of plasticity derived in Appendix B (table A1, Appendix A).

For each deme, we recorded the population size, the average non-plastic component, the average plasticity, and the genetic variance every 200 generations. The genotype of each individual was recorded at the end of the simulations and at the end of the burn-in period. We performed 100 independent realisations for each parameter set (unless stated otherwise).

Apart from performing simulations, we analytically estimated plasticity that maximises the mean population fitness locally (i.e., the *optimal plasticity* ; Appendix B). Notably, we derived approximate conditions for when a population with the capacity for plasticity is expected to attain a larger range than a population lacking this capacity.

## 3 Results

### 3.1 Analytical approximation of the optimal plasticity and the critical cost of plasticity

To derive the conditions allowing plasticity to evolve during range expansion over a gradually steepening environmental gradient, we have undertaken the following steps. First, we found that a locally optimal plasticity, *g*_*e*_ (i.e., plasticity that maximises the local mean population growth rate in equilibrium) in a temporally static environments with a given local environmental gradient *b*(*x*) is given by:

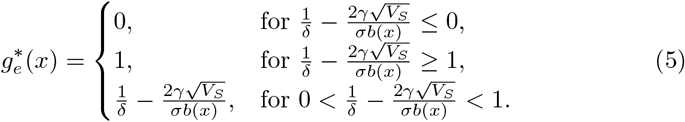

In temporally fluctuating environments, the optimal plasticity is typically larger than in temporally static environments (equation (B41)). Here we explain the implications of the optimal plasticity in the case of static environments, for simplicity, but the same arguments apply to temporally fluctuating environments. Using equation (5), we found a critical environmental gradient (hereafter called the *critical plasticity gradient*), below which the optimal plasticity is zero (i.e. when 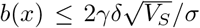). That is, below the critical plasticity gradient any potential positive plasticity that may evolve during initial phases of range expansion is transient, and will eventually vanish.

Next, we made use of the critical plasticity gradient to deduce the conditions allowing a population expanding its range over a gradually steepening gradient to utilise plasticity. Recall that, for a population without the capacity for plasticity, local adaptation is expected to fail at a critical environmental gradient [11] (hereafter *critical genetic gradient*, to emphasise that it corresponds to the case where plasticity is absent). We conclude that when the critical genetic gradient is smaller than the critical plasticity gradient, local adaptation for a population with the capacity for plasticity fails under the same conditions as for a population lacking the capacity for plasticity.

More generally, we show that there are three different regimes for the range margins (figure 1) with respect to two compound parameters, that is *γδ*/*r*_*m*_ and 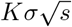 (table 1). The three different regimes are: no difference in the range compared to when the population does not have the capacity for plasticity (this regime, hereafter denoted by *R*_0_, is discussed above and corresponds to the white region in figure 1); a larger, but finite, range than when the population does not have the capacity for plasticity (grey region in figure 1, above dashed line; hereafter denoted by *R*_1_); and potentially infinite range (figure 1, below dashed line; hereafter denoted by *R*_2_).

**Figure 1:**
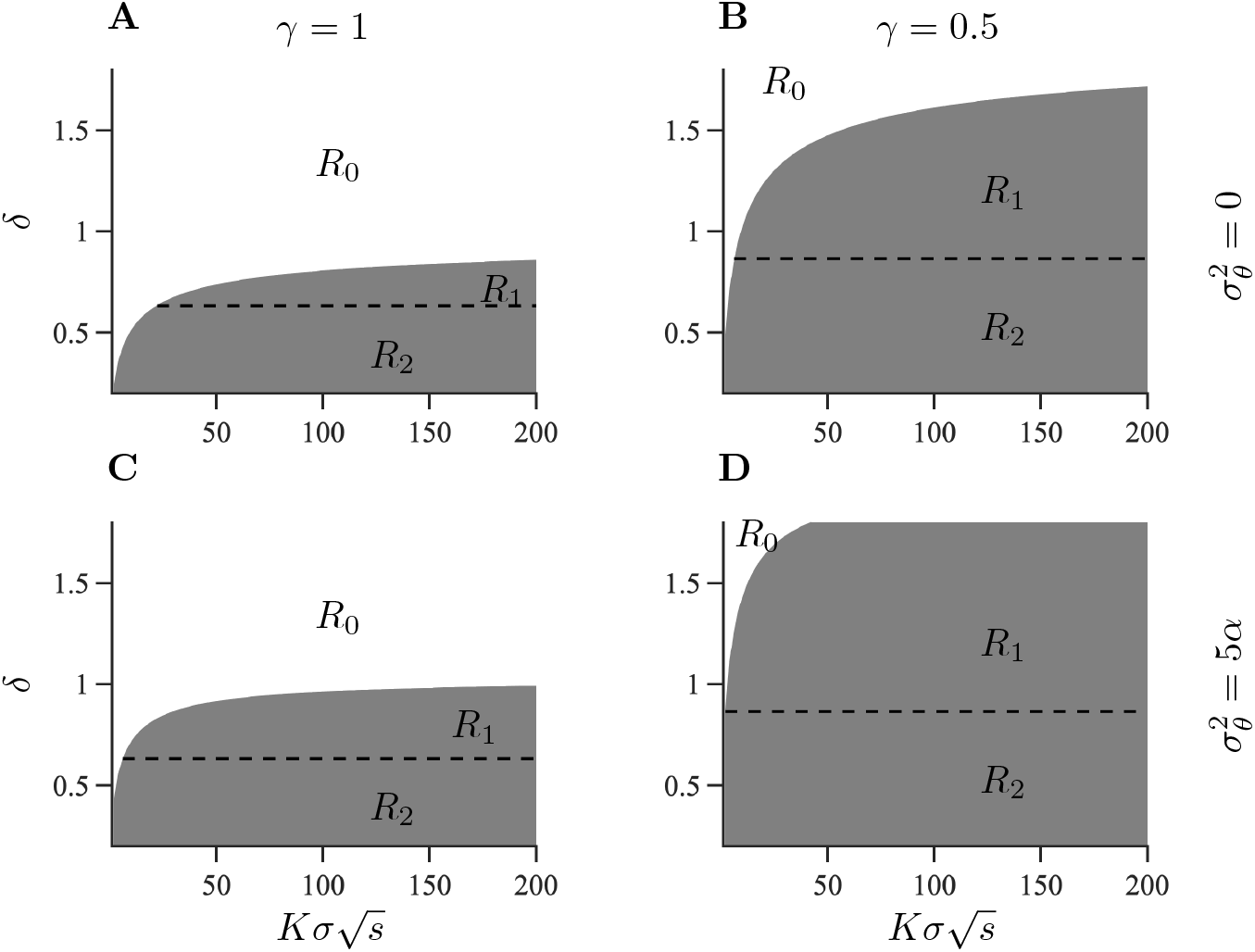
For a given variance of temporal fluctuations in the optimal phenotype, the cost of plasticity divides the parameter space, consisting of the two compound parameters 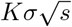 and *γδ*/*r*_*m*_, into three regimes, *R*_0_, *R*_1_ and *R*_2_. In regime *R*_0_ (shown in white), range margins form under the same conditions as without plasticity. In regimes *R*_1_ and *R*_2_ (shown in grey), the range is larger than without plasticity. The dashed line corresponds to a maximum mean population growth rate of zero when the mean phenotype is at the optimum and plasticity equals one. Above the dashed line, in regime *R*_1_, the equilibrium range is finite. In regime *R*_2_ (below the dashed line in the grey area) the growth rate of the population is positive for plasticity of 1. Left column: regimes for a linear cost-related function. Right column: regimes for a concave cost-related function (*γ* = 0.5). Upper row: regimes for a temporally static environment. Lower row: regimes for temporally fluctuating environment where 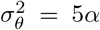 (with 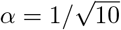).

Finally, we found a critical cost of plasticity (*δ*_*c*_) below which the critical genetic gradient is larger than the critical plasticity gradient. In other words, the critical cost of plasticity is the smallest cost of plasticity for which the dynamics of range expansion fall within regime *R*_0_. The critical cost of plasticity, generalised to account for temporal fluctuations of environmental conditions (Appendix B), is given by

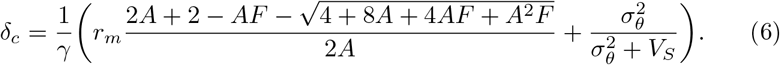

Here, 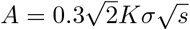 and 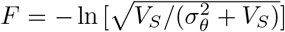 (for notations see table 1). The critical cost (equation (6)), separates the white region from the grey in figure 1.

Outside of the parameter region where regime *R*_0_ is realised, i.e., when the cost of plasticity is lower than the critical cost, the equilibrium range of the population is expected to be larger than for a population without the capacity for plasticity. Here, the equilibrium range is either finite, but larger than for a population without the capacity for plasticity (*R*_1_) or it is possibly infinite (*R*_2_; note that regime *R*_2_ accounts for cases where unlimited ranges occur, but this may not happen for all parameters belonging to regime *R*_2_, as we discuss next).

We distinguished regimes *R*_1_ and *R*_2_ using a necessary but not sufficient condition for unlimited range expansion (dashed line in figure 1), namely that the cost of plasticity is both lower than the critical cost *δ*_*c*_, and sufficiently low to allow a positive population growth rate with plasticity of 1 (hereafter *perfect plasticity* ; Appendix B).

We did not determine the precise conditions allowing unlimited range expansion. However, this is expected at least when there is no cost of plasticity (equations (1)-(3)). We used simulations to examine several parameter sets belonging to regime *R*_2_, focusing on cases with positive plasticity costs.

### 3.2 Simulation results

For comparison, we first ran simulations without plasticity (figure C1). In simulations without plasticity and with static environmental conditions 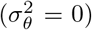, range margins established at the critical genetic gradient (figure C1A), as expected. By contrast, temporal fluctuations in the optimal phenotype (in the absence of plasticity) reduced the range by reducing the equilibrium population size by approximately ln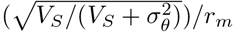 in agreement with [43, 44] (figure C1 B-D; Appendix B). Next, we present simulation results with plasticity.

#### 3.2.1 Temporally static environmental conditions

Recall that our simulations were initialised with a burn-in period. When there were no temporal fluctuations in the environmental conditions, the average plasticity at the end of the burn-in period was close to zero (figure C2). As a consequence, the starting genotype for the non-plastic component was essentially the same as without plasticity (figure C3). Although most alleles for plasticity were fixed, some loci were polymorphic (figure C4).

After the burn-in period, we found that when the cost of plasticity was higher than the critical cost *δ*_*c*_ (so that the expected range expansion dynamics was within regime *R*_0_), plasticity was very low (< 0.05), and the final range agreed with the expected range for populations without the capacity for plasticity (figure 2 A, figures C5 A-B, C6 A-B). This finding was retained when the cost of plasticity was close to the critical cost (figures C5 C and C6 C).

**Figure 2:**
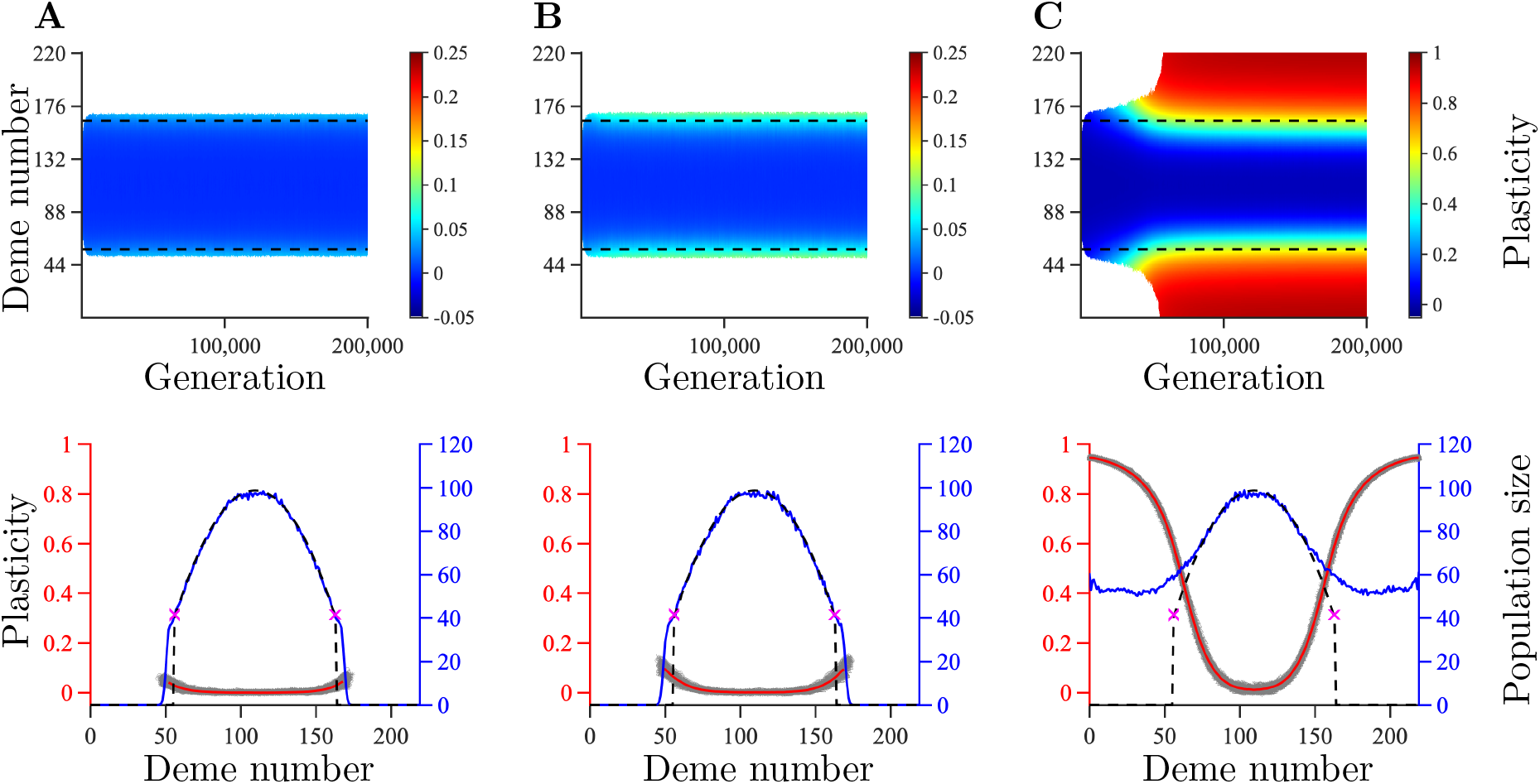
The upper panels show the temporal and spatial evolution of plasticity averaged over 100 realisations during range expansion in a habitat with temporally static environmental conditions. The range expansion dynamics is expected to fall within regime *R*_0_ (column A), *R*_1_ (column B), or *R*_2_ (column C). The columns differ by the parameter *δ*: *δ* = 1.3 (A), *δ* = 0.9 (B), *δ* = 0.5 (C). The red lines in the bottom panels show plasticity averaged over 100 realisations (red axis on the left), the grey areas indicate the spread of plasticity between different realisations. The blue lines show the population size, averaged over 100 realisations (blue axis on the right). The dashed lines in the upper panels denote where adaptation is expected to fail for a population without plasticity. The dashed lines in the lower panels show the expected population size in the absence of plasticity and the purple crosses indicate the expected failure of adaptation. Remaining parameters: *K* = 100, *r*_*m*_ = 1, *V*_*S*_ = 2, *μ* = 10^−6^, *σ* = 1, *L* = 799, *α* = 0.3162, *β* = 0.0013, *γ* = 0.5 and *σ*_*θ*_ = 0.

Conversely, when the cost of plasticity was lower than the critical cost, but sufficiently high to prevent a population with perfect plasticity to have a positive growth rate (i.e., parameters within the expected regime *R*_1_), we observed a higher plasticity in the edges and a slightly larger range than when the cost was above the critical cost (figure 2 B). For a more concave cost-related function, the difference between the ranges attained in regime *R*_0_ and *R*_1_ was larger (compare figure C7 to figure 2 B).

By contrast, when the cost of plasticity was both lower than the critical cost and sufficiently low to allow a population with perfect plasticity to have a positive growth rate, the entire habitat was colonised (figures 2 C, C5 D, C6 D).

Recall that our analytical results (equation (5)) shows that selection favours fully non-plastic (plastic) phenotypes in shallow (steep) environmental gradients. This is in agreement with our simulations (red lines in the bottom panels in figure 2). Regardless of the cost, during the entire simulated time-span, plasticity remained close to zero in the centre of the habitat, where the environmental gradient is shallow. In the edges of the range, plasticity was higher than in the centre of the range. Furthermore, plasticity in the range edges was higher for parameter combination within regime *R*_1_ than for parameter combinations within regime *R*_0_ (average plasticity was 0.02 in figure 2 A, in comparison to 0.1 and 0.7 in figure 2 B and C7 A, respectively). For parameters in regime *R*_2_, the entire habitat was populated and plasticity was close to 1 at the habitat edges (0.95 on average in the case shown in figure 2 C).

The spatial pattern of allele frequencies for the non-plastic component of the phenotype consisted of a series of staggered clines with the same average width as expected for a population without the capacity for plasticity (figure C8). However, when non-zero plasticity evolved, the spacing between the clines was larger than it would have been in the absence of plasticity (e.g. note the absence of clines between deme 10 and deme 50 in figure C8 A, and compare to figure C8 C and E). This is expected by the analogy with [10] (*albeit* in a model without plasticity) because plasticity *g* effectively reduces the environmental gradient by a factor of 1 − *g* [23]. Thus, the spacing between the clines is expected to be increased by a factor of 1/(1 − *g*). For the loci underlying plasticity, allele frequencies increased in a cline-like manner towards the habitat edges for parameters within regime *R*_2_ (figure C8 B) and regime *R*_1_ (figure C8 D). By contrast, when no plasticity evolved, no clear spatial pattern in allele frequencies emerged for the loci underlying plasticity (figure C8 E).

#### 3.2.2 Temporally fluctuating environmental conditions

When the model included temporal fluctuations in the optimal phenotype, results similar to those for static environmental conditions were obtained at the end of the burn-in period when the cost of plasticity was above the critical cost (figure C9 A, B, and D). But, positive plasticity evolved during the burn-in period when the cost of plasticity was low (figure C9 C, E, F, G, H, and I). These results are in agreement with equation (B41) (and see [45]). The spatial patterns of allele frequencies for the non-plastic component at the end of the burn-in period were more noisy than under temporally static environmental conditions (compare figure C10 to figure C3). As for temporally static environmental conditions, the spatial pattern of allele frequencies for the plastic component were irregular (figure C11).

After the burn-in period, when the population was allowed to expand its range, no plasticity evolved when the cost of plasticity was larger than the critical cost (figures 3 A; C12 A, C, and E; C13 A, C, and E), similarly to when the environment was static. In addition, the population size and range extent attained at the end of our simulations were the same as for a population without the capacity for plasticity (compare figure C1 B to figure C13 A; figure C1 C to figure C13 C; and figure C1 D to figure C13 E). Conversely, and similarly to the case with static environmental conditions, when the cost of plasticity was below the critical cost, positive plasticity evolved. For parameters within regime *R*_1_, as expected, the range was larger than in the absence of plasticity, but smaller than the size of the available habitat (figure 3 B). Conversely, for parameters within regime *R*_2_ very high plasticity evolved (on average, 0.95 at the habitat edges in the case shown in figure 3 C) and range expansion continued all the way to the edges of the habitat (figure 3 C; see also figure C12 B, D, F, and figure C13 B, D, F).

**Figure 3:**
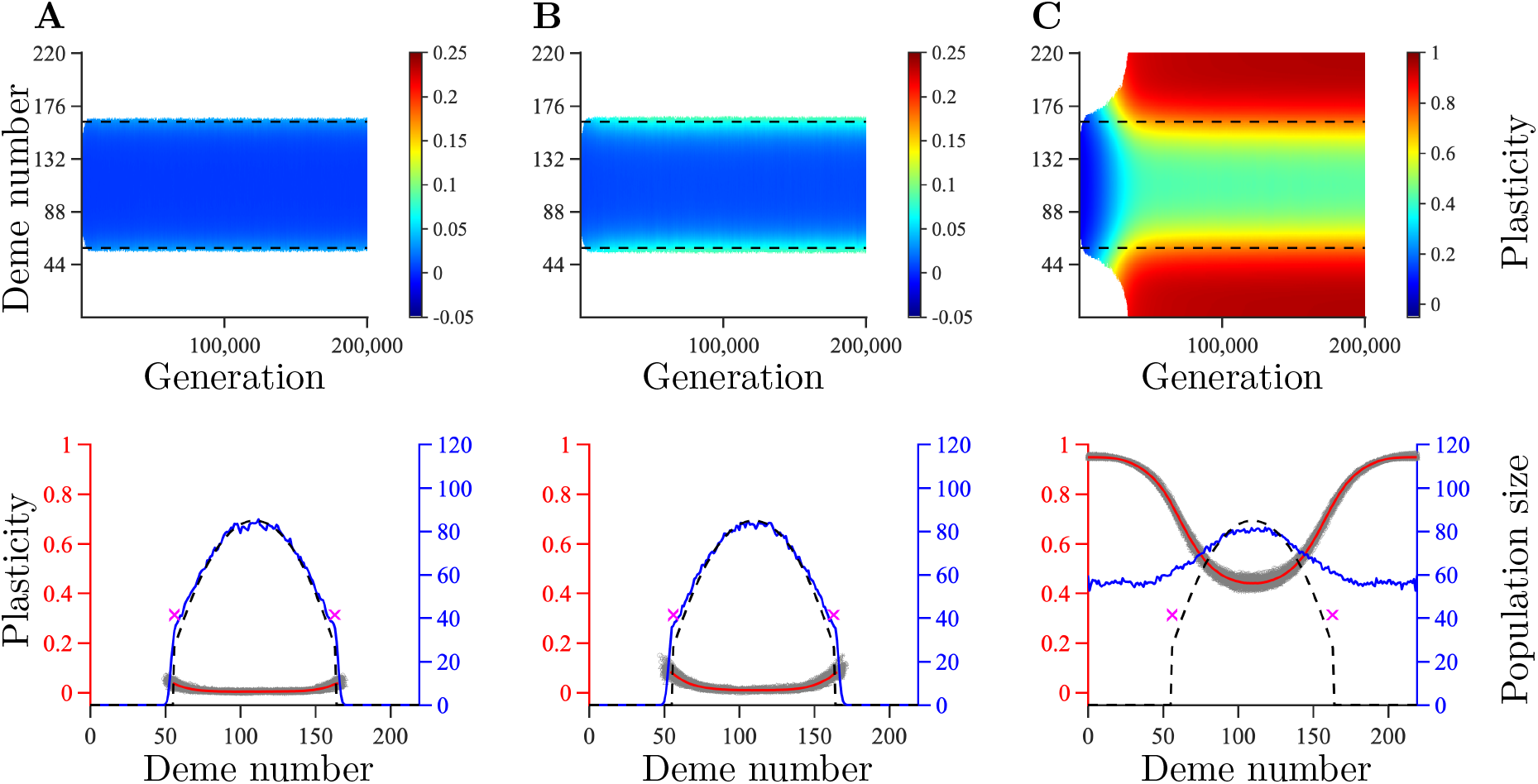
The columns show the results corresponding to those in figure 2 but for temporally fluctuating environmental conditions 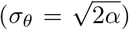. For the parameter values used (apart from *σ*_*θ*_), refer to the caption of figure 2. The dashed lines in <u>th</u>e upper panels denote where adaptation is expected to fail (when 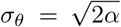) for a population without plasticity. The dashed lines in the lower panels show the expected population size with temporally fluctuating environmental conditions for a population without plasticity. Remaining parameters: *K* = 100, *r*_*m*_ = 1, *V*_*S*_ = 2, *μ* = 10^−6^, *σ* = 1, *L* = 799, *α* = 0.3162, *β* = 0.0013, and *γ* = 0.5.

In contrast to the results with temporally static environments, plasticity in the centre of the habitat was close to zero only when the cost of plasticity was high (red lines in the bottom panels of figure 3 A-B and in figure C13 A, C, E), and it was well above zero in the other cases (red lines in the bottom panel of figure 3 C and in figure C13 B, D, F). Thus, a gradient in plasticity at the end of our simulations was shallower with temporally fluctuating than with temporally static conditions (compare figure 2 C to figure 3 C). Interestingly, at the end of our simulations with temporally fluctuating environmental conditions, plasticity in the centre of the habitat was higher than at the end of the burn-in period (compare, for example, figure C9 C to figure 3 C), and higher than the optimal plasticity given by our approximation (B41). This resulted in a lower population size in the centre of the habitat than the population size expected for a population without plasticity.

## 4 Discussion

Plasticity may facilitate local adaptation to variable and marginal environments, as demonstrated empirically (e.g., [46, 47]), and theoretically (e.g., [9, 23, 24, 25, 35, 38, 48, 45]). However, in some cases the impact of plasticity on local adaptation may be weak or nonexistent (e.g., [15, 26, 32, 49]). The extent to which plasticity is involved in local adaptation may impact on the evolution of species’ ranges and range margins. However, theoretical understanding of the role of plasticity in the establishment of range margins was limited to situations in which genetic variance is an (arbitrarily) fixed, rather than an evolving, property of a population [9] (but see [25]). Importantly, studies of range expansion in the absence of plasticity [8, 10, 11] have shown that genetic variance is a key factor involved in the establishment of range margins. Indeed, fixed genetic variance can cause non-trivial range margins to establish (giving rise to finite ranges, smaller than the size of the available habitat), whereas evolving genetic variance, under otherwise the same model conditions, can allow unlimited range expansion [10]. This suggests that allowing genetic variance to evolve, instead of keeping it fixed, may alter the role of plasticity in the establishment of range margins, both qualitatively and quantitatively. This is the focus of our study. We are primarily interested in situations where populations without plasticity attain non-trivial range margins, such as range expansions over gradually steepening spatial environmental gradients, either without or with temporal fluctuations.

### 4.1 When does the capacity for plasticity increase the range of a population?

Our main result is that plasticity may be involved in the establishment of range margins in one of the following three qualitatively different ways: i) no effect of plasticity, ii) plasticity increases the range by a finite amount, or iii) plasticity allows for unlimited ranges (i.e., absence of non-trivial range margins). Which of these possibilities is realised depends on the benefits of plasticity relative to its costs. Notably, we found a critical cost of plasticity, *δ*_*c*_, above which plasticity does not evolve and the population (despite the capacity for plasticity) is expected to attain the same range as a population lacking the capacity for a plastic response. Below this cost, the range of the population is wider than the range of a population that lacks the capacity for plasticity. Interestingly, the critical plasticity cost is smaller in temporally fluctuating than in static environments, in agreement with [45]. Furthermore, we found a second (smaller) critical cost (hereafter *threshold cost*) below which the range may be infinite (or constrained by a finite habitat size).

When the cost of plasticity is above the critical cost *δ*_*c*_, in local populations up to and beyond the critical genetic gradient (found in [11]), fitness is maximised when plasticity is zero. As a consequence, above the critical plasticity cost, the equilibrium range of a population with the capacity for plasticity coincides with the range of a population lacking this capacity. This is confirmed by our simulation results. Throughout the range, local plasticity was zero on average, except in local populations in the close vicinity of the range margins where slightly positive plasticity evolved. This is expected because marginal populations are demographic sinks (*sensu* [50]). Here, a strongly positive feedback between local maladaptation and small local population size increases local selection for plasticity [9]. Importantly, however, this effect is weak above the critical plasticity cost, making plasticity ineffective to increase the range beyond the range expected in the absence of plasticity.

By contrast, when the cost of plasticity is below *δ*_*c*_, positive plasticity is optimal below the critical genetic gradient. This allows positive plasticity to evolve and be maintained in local populations. In turn, positive plasticity reduces local maladaptation, as well as local selection gradient (as also suggested in [23]), thus making it possible for a population to expand beyond the range expected in the absence of plasticity (i.e., beyond the critical genetic gradient). Interestingly, when the cost of plasticity is so low that the population may simultaneously express perfect plasticity and have a positive growth rate (i.e., below the threshold cost we found), there may be no limit to range expansion (but note that the threshold cost corresponds to a necessary, but not sufficient condition for infinite range expansion to occur). While we were not able to formally prove that infinite range expansion occurs when plasticity costs are sufficiently small, but positive (note that zero costs trivially result in infinite range expansion, as also pointed out in [9], and see references therein), our simulations with non-zero plasticity costs below the threshold cost confirmed that the population occupied the entire habitat (which is necessarily finite in simulations), and that large plasticity evolved (close to 1 at the habitat edges).

Conversely, when the cost of plasticity is below *δ*_*c*_, but still so large that a population with perfect plasticity cannot have a positive growth rate (i.e., above the threshold cost), the capacity for plasticity leads to a range that is finite but larger than when plasticity is absent. Notably, the width of the parameter region where this regime is realised (i.e., between the critical and the threshold cost) is governed by the concavity of the cost function. The more strongly concave the cost function is, the wider is the regime where plasticity leads to finite but larger ranges than when plasticity is absent. For linear or convex cost functions, this regime is very narrow and almost nonexistent for biologically plausible parameters. Consequently, in populations with linear or convex plasticity cost functions, plasticity in equilibrium tends to be either zero throughout the range of the population, or the population may expand its range without limits. We discuss the consequences of this finding in the next subsection.

Recall that we assumed a gradually steepening spatial environmental gradient. Under this assumption, we found a spatial gradient in plasticity when the cost of plasticity was below *δ*_*c*_. This is similar to the pattern found in e.g., [9, 23]. However, in those studies, genetic variance was fixed. Consequently, in [9, 23] the mean population phenotype deviated more from the local optimum further away from the core habitat, resulting in an increased selection for plasticity away from the core habitat. In our model, by contrast, genetic variance is allowed to evolve, meaning that the mean population phenotype in populated areas matches the (average) optimal phenotype. Here, maladaptation is due to genetic variance that increases as the environmental gradient steepens. This increase in genetic variance is further reflected in a progressively decreasing realised population size (although all demes had the same carrying capacity). Thus, in our model, genetic variance increases as the distance from the core population increases, and this results in stronger selection for plasticity. However, we note that the plasticity gradient occurs only below the critical plasticity cost.

Furthermore, in a range-expansion model with environmental conditions that change linearly in space (i.e., with a constant rather than a steepening gradient), and with evolving genetic variance, it was argued that a spatial gradient in plasticity levels out in the long run [25]. To verify this, we performed range expansion simulations along an environment that changes linearly in space (figure C14). We noted a small increase in plasticity towards the habitat edges. This probably reflects edge effects caused by the number of loci we used in simulations (this effect is likely to decrease upon increasing the number of loci, but we did not test this further). However, and as expected, we found that the gradient in plasticity was much shallower when the environmental gradient was constant (figure C14) than when it was steepening (figure 2 C). This is in good agreement with our analysis showing that local plasticity depends on local environmental gradient.

Finally, in our simulations plasticity evolved slower during range expansion than the non-plastic component of the phenotype. This is both due to the steepening environmental gradient, which was shallow in the centre of the habitat, and due to the relatively small allele effect sizes at loci underlying plasticity. By contrast, plasticity evolved much faster in [25], where the environment changed linearly in space and fewer loci were underlying plasticity (so that the allele effect sizes at loci underlying plasticity were larger). Indeed, in our simulations with larger allele effect sizes at loci underlying plasticity (figure C15), or with a constant, rather than steepening, environmental gradient (figure C14), plasticity evolved faster.

### 4.2 Plasticity costs: empirical data and a lesson from theory

We have analytically re-derived the theoretically well-known result that in the absence of costs, perfect plasticity will eventually evolve [9, 51], and the population would be able to expand its range infinitely. The existence of finite ranges even in the absence of any evident geographical barriers [6, 7], thus, suggests that some limits or costs of plasticity may be involved [33]. However, empirical evidence for plasticity costs have so far been elusive [34, 52, 53], except for a few special cases, such as learning-ability [53]. Our results imply that finding empirical evidence for plasticity costs may be specifically difficult when cost functions are much more sensitive to high values of plasticity than to low values (i.e., when cost functions are concave). This is because plasticity would be only weakly costly when plasticity is low or moderate. However, plasticity would still be limited, because high plasticity would exert high costs potentially causing a local population to shrink in size (see discussion above). Thus, concave cost functions of plasticity may potentially limit plasticity while rendering costs difficult to detect. Based on this, we speculate that plasticity costs are more likely to be concave than convex in natural populations, but this is yet to be formally demonstrated.

We note that our results are based on the assumption that the cost of plasticity is constant over space and time. If plasticity costs can evolve, they may decrease over time. However, whether the costs of plasticity will eventually vanish remains an open question for future work.

### 4.3 Limitations of the model

The impact of plasticity on local adaptation may be limited by unreliable environmental cues [54, 55, 56]. Because plasticity may be expressed during different life stages of an organism [57], a mismatch between the environment experienced during development of the plastic response and the environment experienced during selection can occur [33]. In this case, high plasticity during the juvenile life stage may produce a population that is overfitted to the temporal environment, and hence ill adapted to future fluctuations in the environmental conditions. It has been shown both theoretically [38, 45, 48, 54, 55, 56, 58] and empirically [59] that this may impede the evolution of plasticity. Note, however, that the expression of plasticity may occur once during a short critical life-stage or reversibly throughout the life of an individual [60, 61]. The cost of unpredictable cues may be less pronounced for reversible plasticity (compared to when plasticity is irreversible), but this depends on the cost for producing the plastic responses, if such costs are present [58]. In our model, we assumed that the environment of development was perceived without noise and that it was the same as the environment of selection. We leave for future studies to investigate how unreliable cues contribute to the formation of range margins.

Recall that we assumed that all loci recombine freely. Thus, we did not explore the effect of reduced recombination between the loci underlying the non-plastic and/or the plastic component of the phenotype. Dispersal in a spatially heterogeneous environment generates linkage disequilibria between loci, which may lead to maladaptive associations between alleles. This may, in turn, promote the evolution of increased recombination [62]. However, the opposite may be true in marginal habitats [40, 63, 64]. Indeed, locally beneficial combinations of alleles may be partially protected from maladaptive gene flow if the recombination rate between adaptive loci is low. This may allow populations to persist along environmental gradients steeper than the critical genetic gradient [40]. Reaching gradients above the critical genetic gradient may allow the population to evolve plasticity even when its cost is above the critical cost. Hence, reduced recombination may potentially allow the evolution of higher plasticity in the range margins than when recombination between the adaptive loci is free. However, reduced recombination between loci underlying plasticity and loci underlying non-plastic local genetic adaptation may cause trade-offs that limit the utility of plasticity [65]. Additionally, reduced recombination may possibly lead to more frequent evolution of maladaptive plasticity due to poor purging of alleles coding for maladaptive plasticity. We leave for further studies to investigate the role of recombination in the evolution of plasticity, and how recombination and plasticity interact to form range margins.

### 4.4 Applications to conservation

It is well-known that ongoing global climate change is expected to cause directional changes in environmental conditions [66]. However, climate change may also be reflected in stronger temporal fluctuations of environmental conditions in many areas [67]. Management and conservation efforts aimed at mitigating the impact of global climate change should therefore include knowledge and predictions on how temporal fluctuations affect the evolution of natural populations. Specifically, we found that unpredictable conditions may lead to decreased ranges of populations that lack the capacity for plasticity for the trait under selection, or for which the capacity for plasticity in the trait under selection is too costly. By contrast, the ranges of populations that have capacity for plasticity with a sufficiently low cost may not suffer any adverse effect from environmental fluctuations (unless the correlation between the environment of development and the environment of selection is weak, as discussed above, or the fluctuations are so strong that the population goes extinct before it can evolve sufficient plasticity). Indeed, temporal fluctuations may promote the evolution of plasticity to such an extent that future range expansion may be facilitated in comparison to when the environmental conditions are static. This is only true, however, when the cost of plasticity is sufficiently low, as our results show. More generally, our results show how the key parameters, including the carrying capacity, the maximal intrinsic growth rate, and plasticity costs, jointly impact on the conditions a population may adapt to and tolerate. Notably, we show that enhancing the growth rate or the carrying capacity of a population may potentially facilitate the evolution of plasticity and thereby increase the range of conditions a population may endure, We, therefore, suggest that the parameters identified in our analytical treatment, notably the carrying capacity, the maximal intrinsic growth rate, and plasticity costs should be taken into account, for example, when designing assisted evolution programmes aimed at increasing the tolerance of populations to future climate change [68, 69, 70, 71].

Furthermore, invasive species are a major threat to biodiversity worldwide [72]. Invasive species often exhibit higher plasticity than non-invasive species do [73, 74, 75] and it has been suggested that plasticity may be a key factor governing the invasion success of invasive species [73, 75, 76]. Here, we emphasise that a key factor may, instead, be the cost of plasticity for the trait under selection relative to the critical cost of plasticity. Thus, management of ecosystems aimed at preventing the spread of invasive species should take plasticity and, specifically, the critical cost of plasticity into account [77, 78, 79]. This will be particularly important for mitigating potentially elevated risks of biological invasions associated with climate change [77, 80].

### 4.5 Conclusion

We identified the key parameters that determine when the capacity for plasticity increases the range of a population. Specifically, we derived an approximation for the critical plasticity cost above which plasticity is detrimental to the population throughout its entire range. Our results suggest an important role of plasticity costs for range expansions and persistence of ranges, not least in the face of increasingly temporally unstable environmental conditions.

## 5 Authors’ contributions

Conceptualization: MR. Data curation: ME and MR. Formal analysis: ME and MR. Funding acquisition: MR. Investigation: ME and MR. Methodology: ME and MR. Project administration: ME and MR. Resources: MR. Software: ME. Supervision: MR. Validation: ME and MR. Visualization: ME and MR. Writing – original draft: ME and MR. Writing – review & editing: ME and MR.

## 8 Acknowledgements

We thank all the speakers and participants of the Webinar “Evolution of Species Ranges” held online March 8-10, 2021, for their valuable comments and discussions on the topic.

## APPENDIX A Additional model details

In this appendix, we present additional details regarding the individual-based model outlined in the main text.

## Optimal phenotype

The habitat was modelled as a one-dimensional chain of *M* = 220 demes. The mean optimal phenotype in deme *i*, denoted by 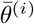, was assumed to be a cubic polynomial of *i*

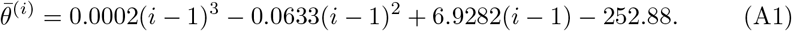

The coefficients in equation (A1) were chosen in such a way that the polynomial had an inflection point in the centre of the habitat, and 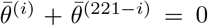 for *i* = 1, 2, …, 220. Furthermore, the gradients in the edges were chosen to be sufficiently steep to make sure that range margins would form well before the edges for a population without plasticity. We allowed the environmental conditions to fluctuate in time in such a way that the optimal phenotype in deme *i* in generation *τ* was

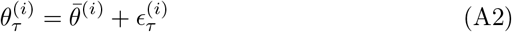

where 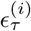 is a random number sampled from the normal distribution with mean 0 and standard deviation *σ*_*θ*_, independently for different demes and at different generations. Table A1 lists the values of *σ*_*θ*_ that we explored.

Note that the coefficients in equation (A1) are different from the coefficients in equation (A1) in [40]. In particular, the habitat in this study contains more demes, and the steepness of the gradient increases faster than in [40]. This was done for practical reasons, to allow for potentially high plasticity to evolve before the population reaches the habitat edges.

For comparison, we also performed a set of simulations of range expansion along an environment that changes linearly in space (without temporal fluctuations). The constant gradient was chosen according to the following two criteria. First, the gradient should be steep enough to allow high plasticity (equation (5) in the main text) to be obtained with plasticity cost parameters *γ* = *δ* = 0.5. Second, the gradient should not be so steep that global extinction is expected to occur, that is, the gradient must be smaller than the critical genetic gradient (defined in the main text; see also [11]). To satisfy these criteria, the phenotypic optimum was chosen to be 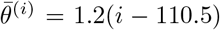. Note that this constant gradient is less steep than the gradient required to attain the phenotypic optima in the edges of the habitat realised in our model (equation A1) with a steepening gradient (the phenotypic optima in the edges for the above chosen constant gradient are ±131.4 in contrast to ±252.9 for the steepening gradient).

In the simulations where the environmental gradient was constant, the allele effect sizes were chosen to ensure that the selection per allele in the edges of the habitat was the same as in the main model for both the non-plastic and the plastic component of the phenotype. To satisfy this requirement, we kept the allele effect sizes for the non-plastic component of the phenotype the same as in the main model (with a steepening gradient), while the allele effect sizes for the plastic component were chosen in such a way that the expressed plasticity per allele was the same in the habitat edges as for the main model (that is, the effect sizes of single alleles were increased by a factor 252.9/131.4 whereas the phenotypic optimum in the edges was decreased by the same factor, thus keeping the plastic response per allele constant). This, in turn, allowed us to employ a smaller number of loci in simulations where the environmental gradient was constant (L=415) as compared to the simulations where it was steepening (*L* = 799). Thus, if all loci underlying the non-plastic component were homozygous for alleles with effect size *α/*2 in deme *M* (or homozygous for alleles with effect size −*α/*2 in deme 1), the edge populations would be perfectly adapted to the local environmental conditions. Similarly, the maximal plastic response was chosen to be ±2, as in the main model. The remaining model parameters were the same as those employed in the main model.

**Figure A1:**
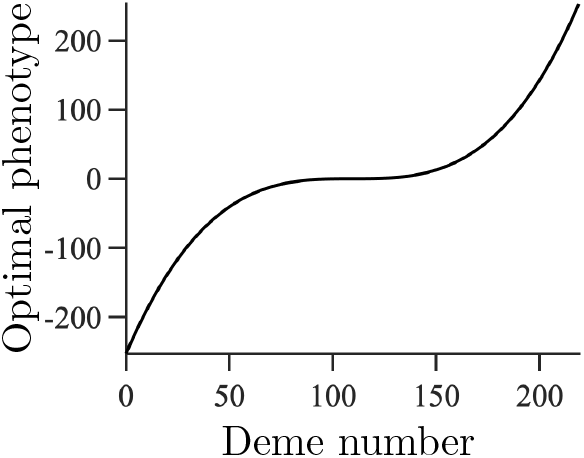
Optimal phenotype as a function of deme number. The line corresponds to a symmetric cubic polynomial with a horizontal inflection point in the centre of the habitat. For this polynomial, the optimal phenotype ranges between −252.9 in the leftmost deme to +252.9 in the rightmost deme.

## Initialisation of simulations

At the start of each simulation, the population occupied *M/*5 = 44 adjacent demes arranged side-by side around the centre of the habitat. The starting genotypes were generated in the following way. For the non-plastic component of the phenotype, 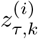, we used the approach explained in [40]: 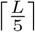 loci (where [*y*] denotes the smallest integer larger than or equal to *y*) were chosen at random and assigned allele frequencies according to the clines at migration-selection equilibrium [10]. Among the remaining loci, half were chosen uniformly at random to be homozygous for alleles with effect size 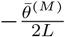, and the remaining loci were chosen to be homozygous for alleles with effect size 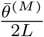 (if the number of remaining loci was odd, one more locus was chosen to be homozygous for the allele with effect size 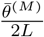). The same loci were chosen to be homozygous for the same allele for all individuals and in all demes. Thus, the average phenotype of the population followed the local optimum initially.

Conversely, for plasticity, one half of the loci were chosen uniformly at random to be homozygous for alleles of effect size 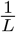 and the remaining loci were chosen to be homozygous for alleles of effect size 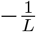 (when the number of loci was odd, one randomly chosen locus was chosen to be heterozyogous). As for the non-plastic component, the same loci were chosen to be homozygous for the same allele for all individuals and in all demes. Thus, plasticity was initially set to zero for all individuals within the starting population, and the genetic variation for the plastic component of the phenotype was minimised throughout the initially occupied habitat.

## Cost-related function

We assumed in the model that plasticity may be more or less costly, with the cost determined by a function 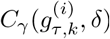 of the form

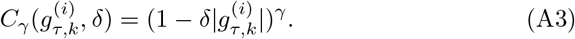

Here, 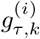 denotes plasticity for individual *k* in deme *i*, in generation *τ*, and the parameters *δ* and *γ* are non-negative parameters, assumed to be constant over time and the same for all individuals (see Methods in the main text for more details). The effect of 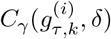 on the fitness of individuals with costly plasticity, in comparison to the fitness of individuals without any cost of plasticity (i.e., when 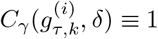) is illustrated graphically in Fig A2.

**Figure A2:**
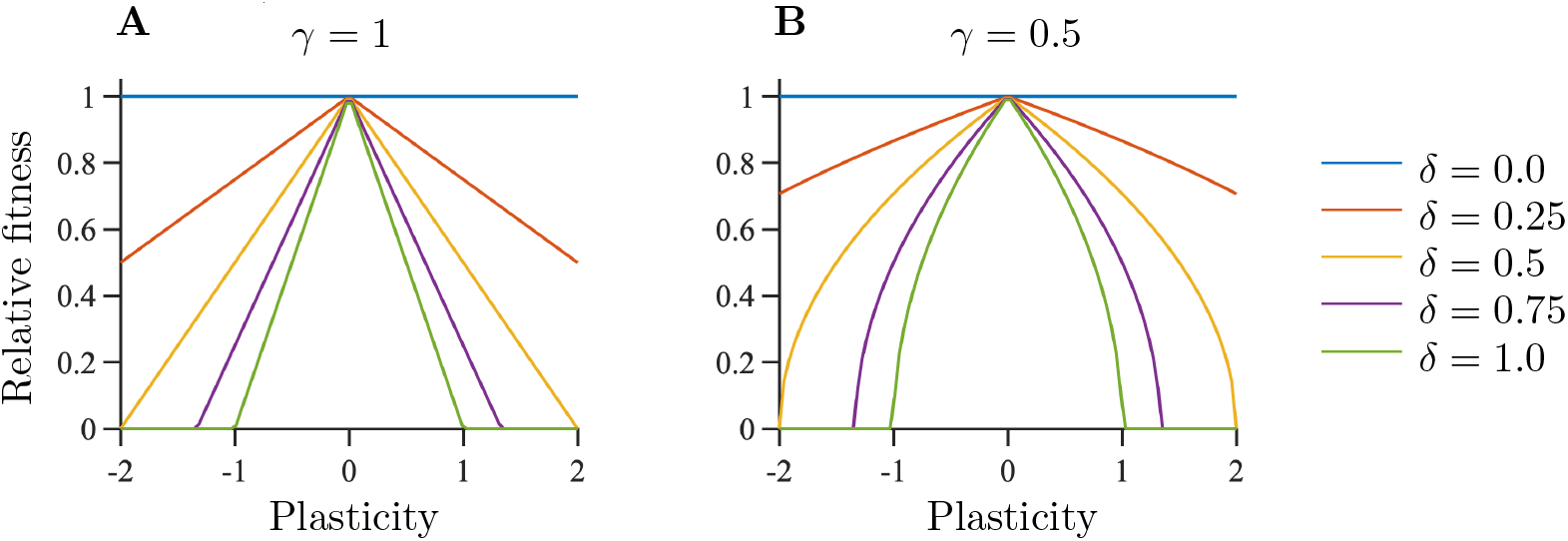
Reduction in fitness due to the cost-related function, 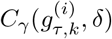. A value of 1 in this figure indicates that there is no cost of plasticity and a value of 0 indicates that the cost is so high that the individual does not reproduce at all (it has a fitness of zero). The lines intersect the *x*-axis when 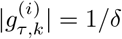.

## Allele effect sizes at loci underlying plasticity

Note that *α* is approximately 126 times larger than *β* (table A1; but we also ran simulations with larger *β*, figure C15). The relative difference between the effects that the two kinds of alleles have on the phenotype differs depending on 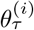 (i.e., the distance from the reference environment). In the habitat edges, the average contribution to the phenotype from an allele with effect size +*β/*2 underlying plasticity is 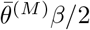, which is twice as large as *α/*2. By contrast, the contribution to the phenotype from an allele with effect size +*β/*2 coding for plasticity approaches zero in the centre of the habitat. The contribution from any allele coding for the non-plastic component is equal to the effect size of that allele, independently of deme position.

## Parameter choices

As explained in the main text, analytical calculations based on a simplified model suggest three qualitatively different regimes of the realised population dynamics with respect to the expected range and the plasticity throughout the range. Which of these three regimes is realised depends on the following parameters: the cost of plasticity, governed by a scale parameter (*δ*) and a shape parameter (*γ*) relative to the maximal intrinsic growth rate, *γ*/*r*_*m*_; the parameter 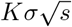 (i.e., the strength of selection per locus on the non-plastic component of the phenotype multiplied by the maximum local population density); the parameter 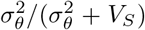 (i.e., the variance of the temporal fluctuations in the optimal phenotype relative to 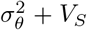; and the shape of the function that determines the optimal phenotype, 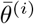 (see table 1 in the main text for a list of parameters). The three different regimes are: 1) no difference in the expected range compared to when the trait under selection is purely non-plastic (denoted by *R*_0_ throughout), 2) a larger, but finite, range for populations with capacity for plasticity, compared to when the trait under selection is purely non-plastic (denoted by *R*_1_ throughout), and 3) a regime where infinite range expansion may occur (denoted by *R*_2_ throughout). An illustration of where in the parameter space these three regimes are realised is shown in figure 1. Further details are given in Appendix B.

To test the main predictions from the above-mentioned analytical calculations, and to confirm the existence of these three regimes, we used computer simulations for different values of the first three among the four parameters mentioned above (i.e., we varied *δ*, *γ*/*r*_*m*_ (keeping *r*_*m*_ constant), 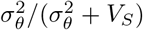 (keeping *V*_*S*_ constant), and 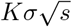 (keeping *σ* and *s* constant), but we did not vary 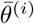). The parameters were chosen to be within each of the three possible regimes, and we included a few borderline cases to test for robustness of the analytical results.

First, for the regime *R*_0_ and no temporal fluctuations in the optimal phenotype, we used cost parameters *δ* = 0.75 or *δ* = 0.6 for *γ*/*r*_*m*_ = 1 (*δ* = 0.6 being very close to regime *R*_1_ and/or *R*_2_) and *δ* = 1.3 for *γ*/*r*_*m*_ = 0.5, with 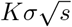 set to 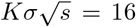 in each case. To further validate our results, we assessed the model outcomes with a parameter combination where the cost of plasticity was sufficiently low to allow the population to have a positive growth rate if the phenotype would be determined entirely by plasticity although equation (6) predicts the range margin to be established at the critical genetic gradient. This parameter combination was 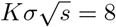, *δ* = 0.5 and *γ*/*r*_*m*_ = 0.5. Notably, in this case our simulation results indeed show that the realised dynamics fall within regime *R*_0_, in line with our analytical results. This is because, as we show in Appendix B, the critical gradient *sensu* [11] is shallower than the smallest gradient where non-zero plasticity improves the population’s mean fitness and hence plasticity either does not evolve at all, or it is not maintained in the long run. For regime *R*_0_ with temporal fluctuations in the optimal phenotype, we used 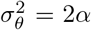 and the cost parameters *δ* = 0.75 for *γ*/*r*_*m*_ = 1 and *δ* = 1.3 for *γ*/*r*_*m*_ = 0.5. In both cases we used 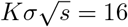.

**Table A1:**
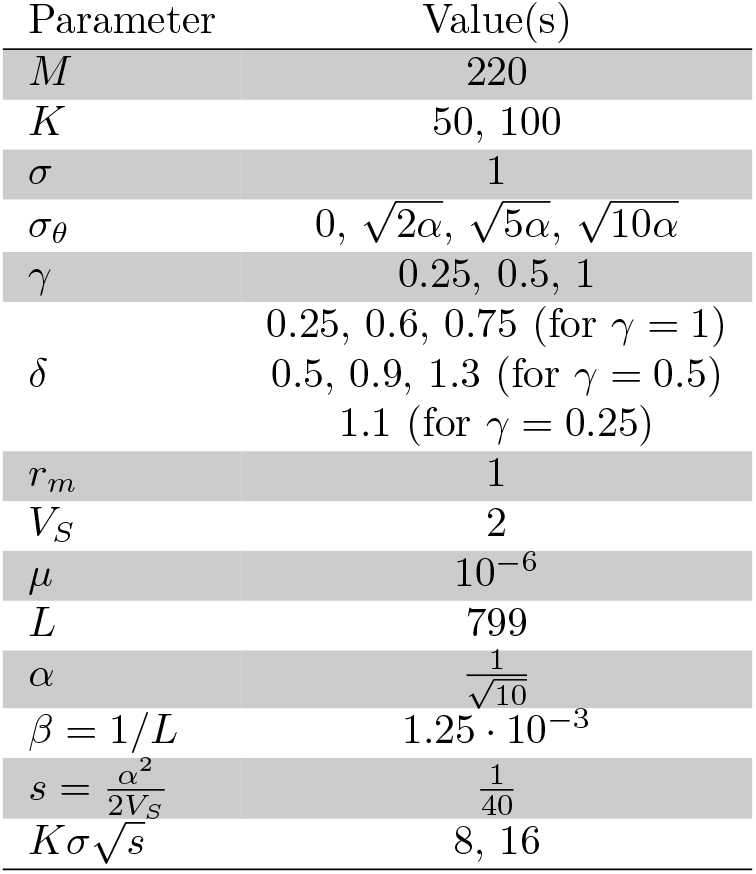
Parameter values examined. Note: Among the parameter combinations listed, we performed simulations with a subset of combinations, allowing us to capture qualitatively similar as well as different simulation outcomes.

Second, for the regime *R*_1_ and no temporal fluctuations in the optimal phenotype, we used *δ* = 0.9, *γ*/*r*_*m*_ = 0.5 and *δ* = 1.1, *γ*/*r*_*m*_ = 0.25. In both cases, we used 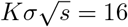. For regime *R*_1_ with temporal fluctuations in the optimal phenotype, we used the following parameter combinations: 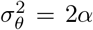 with the cost parameters *δ* = 0.9 and *γ*/*r*_*m*_ = 0.5; 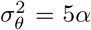 or 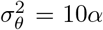, with the cost parameters *δ* = 0.75 and *γ*/*r*_*m*_ = 1,*δ* = 0.9 and *γ*/*r*_*m*_ = 0.5, or *δ* = 1.3 and *γ*/*r*_*m*_ = 0.5. In all cases we used 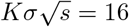.

Third, for the regime *R*_2_ and no temporal fluctuations in the optimal phenotype, we used *δ* = 0.25 for *γ*/*r*_*m*_ = 1 and *δ* = 0.5 for *γ*/*r*_*m*_ = 0.5. In both cases, we used 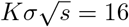. For regime *R*_2_ with temporal fluctuations in the optimal phenotype, we used the following parameter combinations: 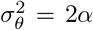 with the cost parameters *δ* = 0.25 and *γ*/*r*_*m*_ = 1, or *δ* = 0.5 and *γ*/*r*_*m*_ = 0.5; 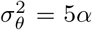 or 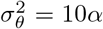, with the cost parameters *δ* = 0.25 and *γ*/*r*_*m*_ = 1, or *δ* = 0.5 and *γ*/*r*_*m*_ = 0.5. In all cases we used 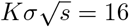.

For each regime, we ran 200, 000 generations for the parameters with *γ* = 0.5 when 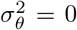 or 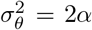, or when *γ* = 0.25, and 100, 000 generations for the remaining parameter combinations. The results are shown and discussed in the main text (see also Appendix B).

## Appendix B Analytical approximations for the optimal plasticity

In this appendix, we derive an analytical approximation for the plasticity that locally maximises the average population growth rate in equilibrium (hereafter called the *optimal plasticity*). This approximation gives an estimate for when the capacity for plasticity cannot increase the equilibrium range of the population, compared to the expected range of a population that does not have the capacity for plasticity. Although the approximation relies on many simplifying assumptions, it gives a qualitatively good agreement with our simulation results.

In what follows, we describe the model we use to carry out the analytical calculations in this appendix. This model is a simplified version of the model described in the main text. The model simplifications here were made to ease the analytical treatment of the system. In this appendix, we assume that the local population density is so large that drift can be neglected (but this assumption is relaxed in subsection B4), and perform the derivations in a continuous onedimensional space, *x*, and continuous time, *t*. The discrete case can be obtained by defining Δ*x* as the distance between two neighbouring demes and Δ*t* as the time between two successive generations. Individuals are assumed to be diploid. We use *θ*(*x, t*) to denote the optimal phenotype in position *x* in time *t*. As in [10], we assume that the optimal phenotype can be approximated locally by a function that changes linearly in space. We denote by *z*(*x, t*) the non-plastic component of the phenotype of an individual, considered as an observation of a random variable sampled over the population in position *x* and time *t*. We assume that plasticity is constant in time and, locally, in space, and we denote it by *g*. Recall that the phenotype, denoted by *u*(*x, t*), is the sum of a nonplastic component, *z*(*x, t*), and a plastic component, *gθ*(*x, t*) (the latter being the product of plasticity *g* and the optimal phenotype *θ*(*x, t*) of the trait under selection), that is:

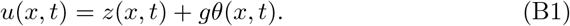

We assume that the non-plastic component is underlain by *L* freely recombining loci, and that there are two possible effect sizes for alleles coding for the nonplastic component, i.e., ±*α/*2. We use 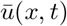 and 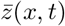 to denote the expected value of the population mean in position *x* and time *t* for the phenotype and the non-plastic component of the phenotype, respectively (hereafter called the *mean phenotype* and the *mean non-plastic component of the phenotype*, respectively). For a given *g* and a given population variance *V*_*z*_ of the non-plastic component of the phenotype, the rate of change of the mean phenotype and of the local population density are given by [81]

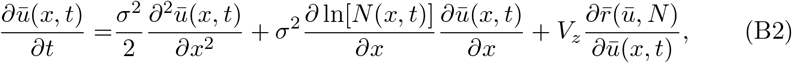

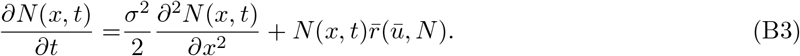

Furthermore, in linkage equilibrium, the rate of change of the allele frequencies *p*_*z,j*_ at locus *j* are given by [10]

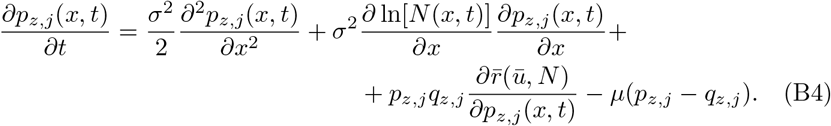

Here, *μ* denotes the mutation rate and *q*_*z,j*_ = 1 − *p*_*z,j*_. We define the continuous growth rate for an individual with phenotype *u*(*x, t*) and plasticity *g* in a location where the optimal phenotype is *θ*(*x, t*) and the population density is *N* (*x, t*) as

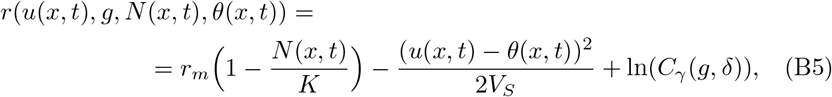

where *r*_*m*_ denotes the maximal intrinsic growth rate, *K* denotes the local carrying capacity, and *V*_*S*_ denotes the width of stabilising selection. Note that *r*(*u*(*x, t*), *g, N* (*x, t*), *θ*(*x, t*)) depends on *z*(*x, t*) through *u*(*x, t*), *g* and *θ*(*x, t*). The function

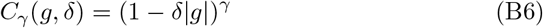

denotes a cost-related function for plasticity, where the non-negative parameters *γ* and *δ* determine the degree of convexity/concavity of the cost-related function, and the threshold plasticity above which the maximal fitness of an individual is zero ( |*g*| < 1*/δ*), respectively (see Methods for a more detailed description). Note that the growth rate, given by equation (B5), corresponds to the logarithm of the discrete fitness function divided by two, i.e. (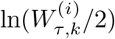) (see equation (2) in the main text).

As mentioned above, we use this simplified model to find the optimal plasticity of the population. The derivation is explained next.

## B.1 Optimal plasticity under static environmental conditions

To find the optimal plasticity, we first reduce the number of parameters in equation (B5) by re-scaling them. Note that, for locally constant plasticity, the deviation of phenotype *u*(*x, t*) from the local optimum can be expressed in terms of the non-plastic component of the phenotype (*z*(*x, t*)) and a re-scaled optimum ((1 − *g*)*θ*(*x, t*)) [23]

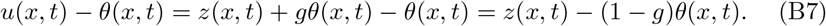

Moreover, for any general cost-related function (including the one considered in our model), which we denote here by *c*_*g*_ (for simplicity and to emphasise the generality) the following holds

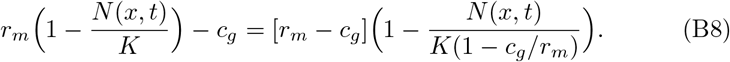

Thus, upon re-scaling the parameters *θ*(*x, t*), *r*_*m*_ and *K* as follows

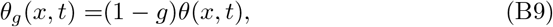

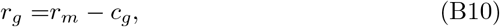

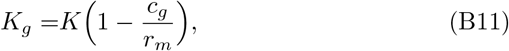

we re-write equation (B5) as

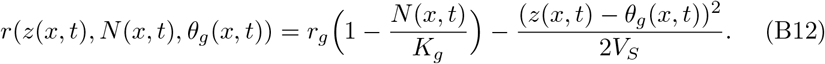

For the remainder of this subsection, we assume that local optimum for the phenotype, *θ*(*x, t*), is kept constant in time. To emphasise the absence of temporal fluctuations in the environmental conditions in the following calculations, we use *θ*(*x*) in place of *θ*(*x, t*) and *θ*_*g*_(*x*) in place of *θ*_*g*_(*x, t*). Furthermore, recall that we assume that both the plastic component of the phenotype and the locally optimal phenotype are determined exactly by the same environmental variable. Under these assumptions we have, thus, reduced the model with constant plasticity to the di-allelic model that has been analysed in [10]. By analogy to [10], it follows that the only stable equilibrium for equations (B2)-(B3) as *t* → ∞ (under the assumption that plasticity *g* is constant) corresponds to a state where the population size locally constant and the average phenotype in position *x* equals *θ*_*g*_(*x*). In addition, the contributions to linkage disequilibrium (LD) from dispersal and stabilising selection in equilibrium cancel out in our model (when selection is weak relative to recombination so that the quasi-linkage equilibrium can be assumed [82]) by the same arguments as in [11].

As stated in the beginning of this appendix, we define the optimal plasticity as the plasticity that maximises the mean growth rate of the population (equation (B12)) in equilibrium. The next step is, thus, to find the population mean of equation (B12) in equilibrium.

We denote the equilibrium population size, and non-plastic component of the phenotype at position *x* by *N*_*e*_(*x*), and *z*_*e*_(*x*). By taking the population mean of equation (B12) in equilibrium, we find

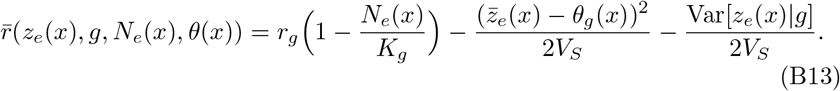

Here, Var[*z*_*e*_(*x*)|*g*] denotes the local migration-selection equilibrium population variance of the non-plastic component of the phenotype *z*_*e*_(*x*) in deme *x*, for given plasticity *g*. Recalling that 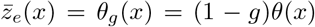 and using the above mentioned analogy to [10], it follows that

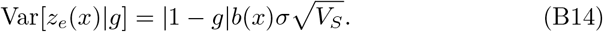

Here, *b*(*x*) = *∂θ*(*x*)/*∂x* denotes the environmental gradient in position *x*.

Upon expressing equation (B13) in terms of *K*, *r*_*m*_, 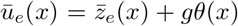, and *c*_*g*_ = *C*_*γ*_ (*g, δ*) = *γ* ln(1 − *δ*|*g*|), and using equation (B14), we find

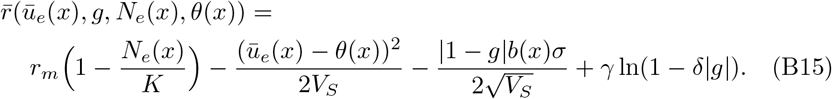

To find the optimal plasticity, denoted by 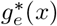, we maximise equation (B15) with respect to *g*, under the requirement that 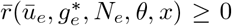, and that 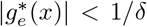 (note that this inequality is strict because the growth rate has singularities at the points where |*g*| = 1*/δ*).

To this end, note that equation (B15) is differentiable when *g* < 0, *g* > 1, or 0 < *g* < 1. In these cases, the derivative of equation (B15) with respect to *g* is given by

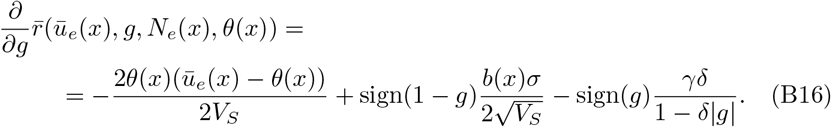

Here, sign(*y*) is the signum function, which is 1 when *y* is positive, −1 when *y* is negative, and it is not defined when *y* = 0. Note that, because |*g*| < 1*/δ* by equation (B6), it follows that equation (B16) is strictly positive for all *g* < 0 and strictly negative for all *g* > 1. Consequently, equation (B15) has a maximum at 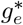 such that 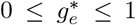 when *δ* ≤ 1, or a maximum such that 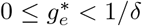 when *δ* ≥ 1 (recall that |*g*| is bounded above by the positive number 1*/δ* according to equation (B6)). The right hand side (RHS) of equation (B16) implies that there is a trade-off between two components of the growth rate when plasticity is increased (assuming 0 ≤ *g* ≤ 1). Increased plasticity decreases the genetic load caused by migration between neighbouring demes (i.e., migration load), which is described by 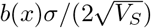 [10], but it increases the cost of plasticity, described by *γδ/*(1 − *δg*). When the benefit of decreasing the migration load is greater than the disadvantage of increasing the cost of plasticity, equation (B16) implies that increased plasticity increases the mean growth rate of the population. Conversely, when the disadvantage of increasing the cost of plasticity is greater than the advantage of decreasing the migration load, equation (B16) implies that decreased plasticity increases the mean growth rate of the population.

To find the maximum of equation (B15), note that if equation (B16) is strictly negative on the open interval 0 < *g* < *m* where *m* = min (1, 1*/δ*) (the interval is not including the discontinuities that equation (B16) has at 0 and 1 or the singularity it has at 1*/δ*), it follows that equation (B15) has a maximum at 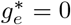. If equation (B16) is strictly positive on the open interval 0 < *g* < 1, it follows that equation (B15) has a maximum at 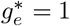.

To find when equation (B16) is strictly positive or strictly negative on the interval 0 < *g* < 1, note that sign(1 − *g*) = sign(*g*) = 1 and |*g*| = *g* when 0 < *g* < 1. Furthermore, the first term in equation (B16) is zero on average in equilibrium because 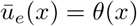. Thus, equation (B16) reduces to

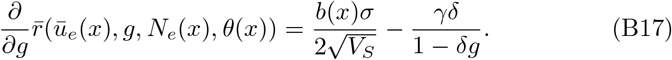

Under the assumption that *g* < 1*/δ*, equation (B17) is a monotonically decreasing function of *g*. Hence, if it is positive for *g* = 1, then it is strictly positive on the interval 0 < *g* < 1. That is, if

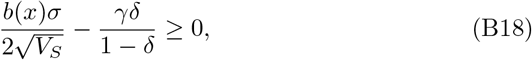

then equation (B17) is strictly positive on the interval 0 < *g* < 1. Similarly, if

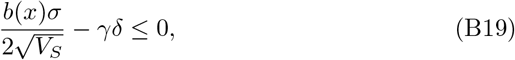

then equation (B17) is strictly negative on the interval 0 < *g* < 1. In other words, when 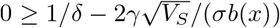, it follows that 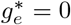. Similarly, when 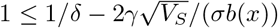, it follows that 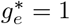.

Otherwise, when 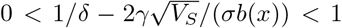, a maximum to equation (B15) satisfies

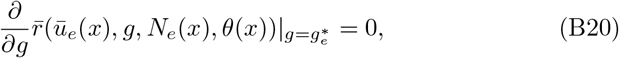

and

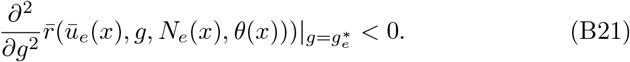

From equation (B20), we find

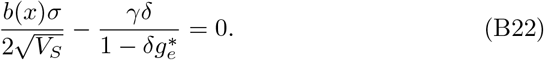

The solution to (B22) with respect to 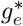 is given by

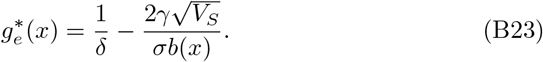

To see that (B23) maximises equation (B15) with respect to *g* in the case when 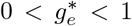, note that the following holds for the second derivative of equation (B15) with respect to *g*, evaluated at 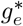.

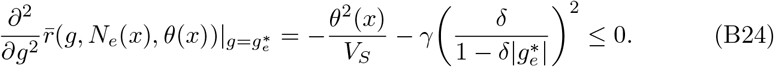

Here, the equality holds if and only if both *θ*(*x*) = 0 and *γδ* = 0. Otherwise, the second derivative is negative. Thus, when at least one of the *θ*(*x*) or *δγ* is nonzero, the second derivative is strictly negative for 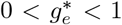. In other words, if 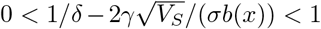, then 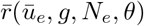 evaluated at 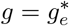, with 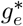 given by equation (B23) corresponds to the global maximum of equation (B15) on the interval 0 < *g* < 1. In the special case when *θ*(*x*) = *δγ* = 0 (i.e., when there is no cost of plasticity and plasticity does not alter the phenotype), the optimal plasticity cannot be defined.

In sum, under the assumptions that at least one of *θ*(*x*) or *δγ* is non-zero, the optimal plasticity, 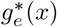, is given by

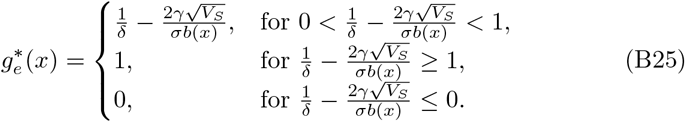

Recall that plasticity effectively reduces the steepness of the environmental gradient, and hence the difference between the phenotypic optima in neighbouring demes (equations (B7), (B9), and (B12); and see also [23]). Consequently, genetic differentiation between local populations is expected to be reduced in comparison to when the trait under selection is not plastic. Equation (B25) implies that when the cost for migration between neighbouring demes 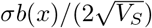 is larger than the cost of plasticity *γδ* (equation (B19)), the population benefits from positive plasticity because the benefit of reducing migration load by 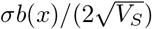 is greater than the cost of reducing the growth rate by *γδ*.

## B.2 Optimal plasticity in a spatially homogeneous environment with temporally fluctuating optimal phenotype

As in the previous subsection, we here aim to find the plasticity that maximises the average fitness of the population (i.e., the *optimal plasticity*). However, in this subsection we allow the optimal phenotype to fluctuate in time, but we assume that the time-average optimal phenotype is constant across space. This allows us to obtain an approximation for plasticity that is expected to evolve during our burn-in simulations (see Methods for details). Note that although the temporal fluctuations in the optimal phenotype were uncorrelated between neighbouring demes in our model, the average effect of the fluctuations over time is the same in each deme. To emphasise the absence of spatial heterogeneity in this subsection, we use *u*(*t*), *N* (*t*), and *θ*(*t*) in place of *u*(*x, t*), *N* (*x, t*), and *θ*(*x, t*), respectively. We will also in this subsection assume that the genetic variance is approximately constant, and denote it by *V*_*G*_. The fitness is, in this case, given by:

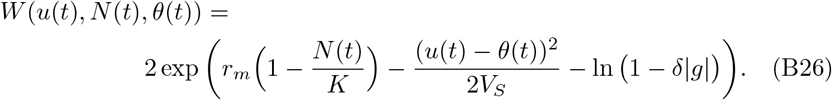

By performing the re-scaling given in equations (B9)-(B11), but with *θ*_*g*_(*t*) in place of *θ*_*g*_(*x, t*), the fitness can be written as

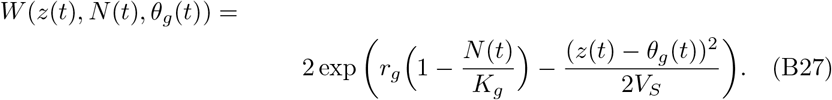

Furthermore, as also pointed out in [83], the variance 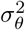 of the re-scaled local optimum *θ*_*g*_(*t*) = (1 − *g*)*θ*(*t*) is given by

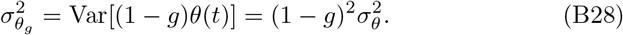

Assuming that the mean population phenotype has a fixed value 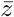 in equilibrium, we find, in accordance with previously published literature (e.g., [43, 44]), that the time-average of the mean population fitness 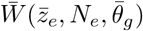 is given by:

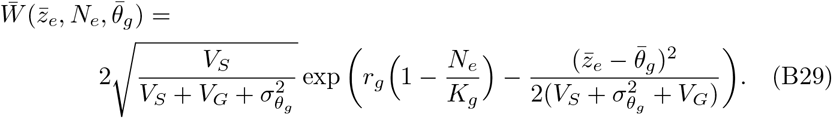

Here 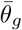 denotes the mean of *θ*_*g*_(*t*). equation (B29) can be rewritten as:

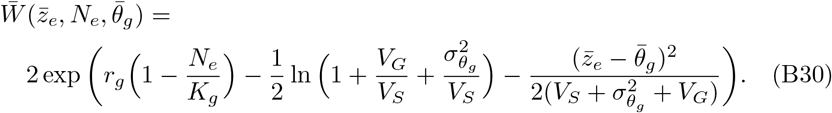

From equation (B30) it follows that the average growth rate of the population is

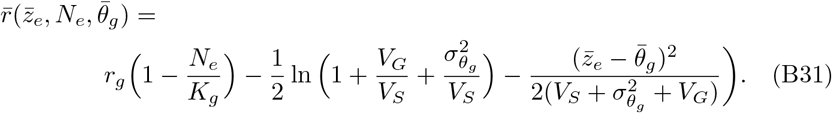

Writing equation (B31) in terms of *u*, 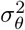, *V*_*S*_, *K*, *r*_*m*_, and *c*_*g*_ = *γ* ln (1 − *δ*|*g*|) yields

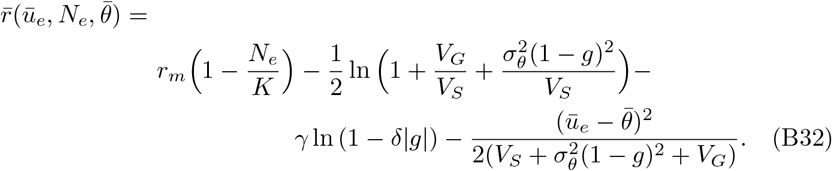

To find the plasticity that maximises equation (B32), we first differentiate equation (B32) with respect to *g*, which in the absence of costs of plasticity:

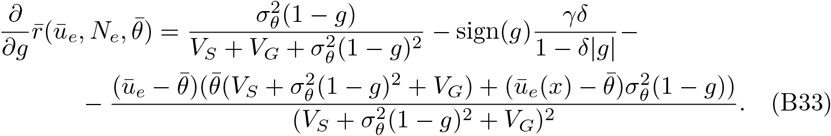

Upon assuming that 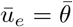 the last term vanishes and equation (B33) reduces to

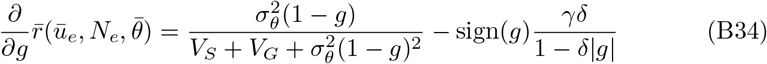

Under the additional assumption that *V*_*G*_ ≪ *V*_*S*_, equation (B34) can be approximated as

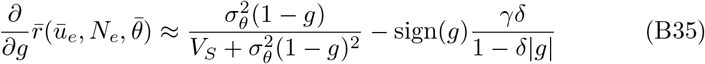

Recall that |*g*| < 1*/δ* (equation (B6)). Therefore, equation (B35) is strictly positive when *g* < 0. Furthermore, note that, under the assumption that *δ* < 1, equation (B35) is strictly negative when *g* ≥ 1 (but approaches zero asymptotically at *g* = 1 as *γ* → 0 or *δ* → 0). Similarly, when *δ* > 1, it follows that equation (B35) approaches −∞ as *g* → 1*/δ*. As a consequence, if equation (B32) has a maximum, this maximum lies on the interval [0, min(1, 1*/δ*)) (note that the interval may include 0 but not 1 or 1*/δ*). To find the maximum of equation (B35) we next assume that 0 ≤ *g* ≤ min(1, 1*/δ*).

For 0 ≤ *g* ≤ min(1, 1*/δ*) equation (B35) is zero when *g* = *g*_0_ such that:

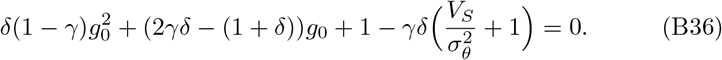

We find:

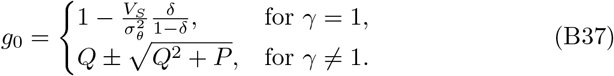

Here, 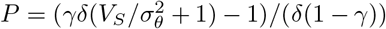, and *Q* = (1 + *δ*(1 − 2*γ*))/(2*δ*(1 − *γ*). To find under which conditions the solutions *g*_0_ to equation (B36) are maxima of equation (B32), we consider the following five different cases with respect to the parameters involved:

## Case 1

*γ* = 1, 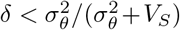. In this case, it follows that 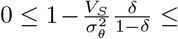 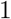, and hence a unique solution to equation (B36) on the interval [0, 1) exists. Furthermore, note that when 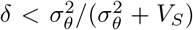, equation (B35) is positive as *g* → 0^+^. Thus, equation (B32) is increasing in a neighbourhood of 0, decreasing in a neighbourhood of 1, it has a unique point, 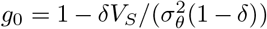, where the derivative is 0 (hereafter referred to as a *critical point*) on the interval [0, 1) and it is continuous. Using the extreme value theorem [84], it follows that *g*_0_ is a maximum.

## Case 2

*γ* ≠ 1, *δ* < 1, 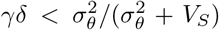. When *δ* < 1, we find that *Q* = (1 + *δ*(1 − 2*γ*))/(2*δ*(1 − *γ*)) ≥ 1. Thus, 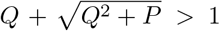, and so 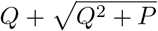 is not a solution to equation (B36) on the interval [0, 1) in this case.

Therefore, there can be at most one solution, *g*_0_, to equation (B36) such that 0 ≤ *g*_0_ ≤ 1. Furthermore, when 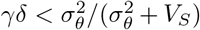 equation (B32) is increasing in a neighbourhood to the right of 0. Because equation (B32) is increasing in a neighbourhood to the right of 0 and decreasing in a neighbourhood to the left of 1, it must attain a maximum, *g*_0_, such that 0 < *g*_0_ < 1. Thus, there is a unique solution, 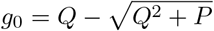, on the interval [0, 1).

## Case 3

*γ* < 1, *δ* ≥ 1, 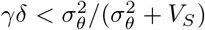. In this case, we first assume that *Q* < 1*/δ*. By re-ordering the terms in *Q*, this assumption can be re-written as:

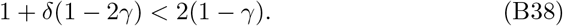

From this, it follows that *δ* < 1. But, because in this case we assume *δ* ≥ 1, it follows that *Q* ≥ 1*/δ*. This further implies that

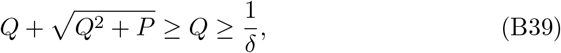

meaning that 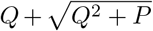 cannot be a solution to equation (B36) for 0 ≤ *g* < 1*/δ*. Thus, there can be at most one solution to equation (B36) for 0 ≤ *g* < 1*/δ*. Furthermore, equation (B32) is increasing in a neighbourhood to the right of *g* = 0 and decreasing as *g* → 1*/δ*^−^, and therefore it must attain a maximum at 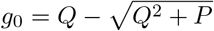.

## Case 4

*γ* ≤ 1, *δ* ≥ 1, 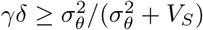. In this case, we find

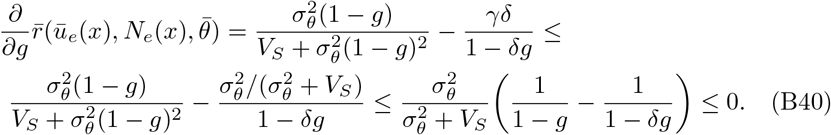

Thus, the derivative is strictly non-positive for 0 ≤ *g* < 1*/δ*. Hence, the maximum of equation (B32) is attained at *g* = 0.

## Case 5

*γ* > 1, *δ* < 1, 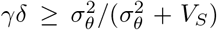. As shown for Case 2 above, when *δ* < 1 it follows that *Q* ≥ 1. As a consequence, equation (B36) can have at most one solution, *g*_0_, within the interval 0 ≤ *g* ≤ 1 (attained when 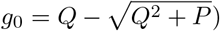). Because equation (B32) is decreasing in a neighbourhood to the right of *g* = 0 and decreasing as *g* → 1^−^ a critical point for 0 ≤ *g* < 1, if it exists, must be an inflection point. Thus, the maximum of equation (B32) is attained when *g* = 0.

Thus, the optimal plasticity under spatially homogeneous but temporally heterogeneous conditions, 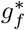, is (cf. equation (B25)):

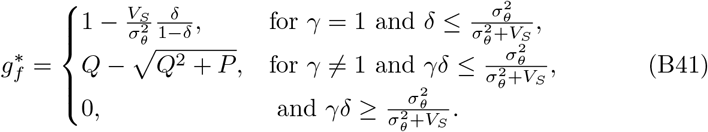

Note that the first- and second-case solutions in equation (B41) are consistent because in the limit of *γ* → 1, the second-case solution converges to the first-case solution, as expected, i.e 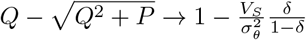 as *γ* → 1.

Equation (B41) was compared to plasticity attained at the end of our burn-in simulations (figures C2 and C9).

## B.3 When is plasticity of zero optimal?

In this subsection, we derive a condition for when the optimal plasticity is zero in an environment where the optimal phenotype changes in space and fluctuates in time. Understanding when plasticity of zero is optimal for a population is of specific interest because, in such cases, the ability to express and evolve plasticity yields no fitness benefit compared to when the capacity for plasticity is absent. The average growth rate of an equilibrium population in a temporally static environment is given by equation (B15). When the optimal phenotype randomly fluctuates in time (but the mean phenotype and the phenotypic variance of the local population are constant over time), the mean growth rate is reduced by the additional load component ln 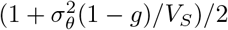 (subsection B2; see also [44]). The mean growth rate, averaged over time, for a population occupying an environmental gradient with temporally fluctuating optimal phenotype is, thus, given by

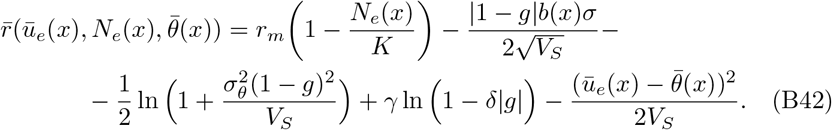

The derivative of equation (B42) with respect to *g* is

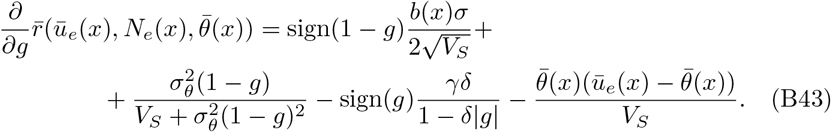

As we found in subsection B2 for the special case when *b*(*x*) = 0, equation (B43) is strictly non-positive on the interval 0 ≤ *g* ≤ min (1, 1*/δ*) when 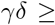 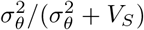. The addition of the term 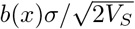, which is independent of *g*, makes equation (B43) strictly non-positive when

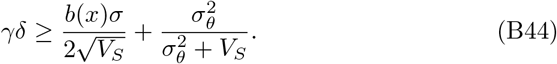

Thus, at any local position *x* where inequality (B44) is satisfied, the mean population growth rate is non-increasing as *g* increases from 0 to min (1, 1*/δ*) (and equation (B43) is zero only at isolated points, otherwise it is zero everywhere, so equation (B42) is decreasing except at isolated points), implying that the local optimal plasticity in such positions is equal to zero. Thus, at any local position *x* where inequality (B44) is satisfied, the local optimal plasticity is zero. Note that, when the fluctuations in the optimal phenotype are small, i.e., when 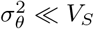, the following holds:

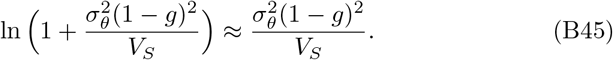

In this case, equation (B44) may be approximated by

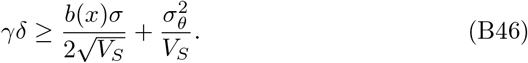

In the next subsection, these results are used to determine the *critical cost* of plasticity, above which the ability to express and evolve plasticity is not expected to facilitate local adaptation anywhere in the range of a population.

## B.4. The effect of plasticity on the critical environmental gradient

In this subsection, the results derived above are used to find an approximate condition for when the ability to express and evolve plasticity increases the range of a population, compared to when plasticity is absent. As shown in [11], a haploid population fails to adapt to the local environment when 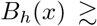 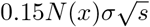 (here, subscript *h* is used to indicate *haploid* populations), where

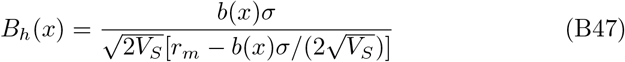

is the *effective environmental gradient*, and 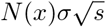 (where *s* = *α*^2^/(2*V*_*S*_)) is the *efficacy of selection relative to drift*. For a diploid population with *N* (*x*) individuals, the population fails to adapt to the local environment when

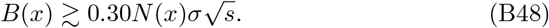

Note that the only difference to [11] is a factor of two in (B48) which accounts for diploidy.

n an environment where the optimal phenotype is temporally fluctuating, the expected population size is reduced in comparison to the expected population size in static environments, due to the load component from the temporal fluctuations, 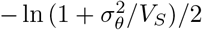 (which may be approximated by 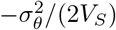 when 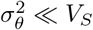. In this case, the population size is given by

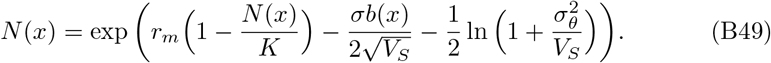

This equation reiterates the results from [44] for the case when 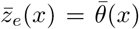. Thus, the composite parameter 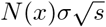 is given by

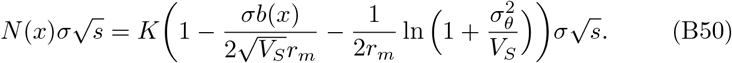

To obtain an expression for the environmental gradient *b*_*c*_ above which local adaptation fails (hereafter the *critical genetic gradient*) for a population without the capacity for plasticity, we write the dimensionless parameters, *B* and 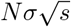, in equation (B48) in terms of the (composite) parameters *b*(*x*), *A*, *E*, and *F*, where

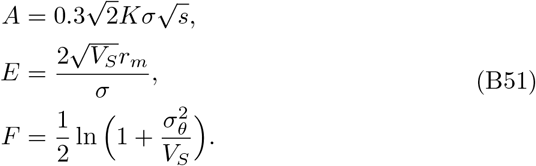

We find that the critical genetic gradient (*b_c_*) for a population without the capacity for plasticity in temporally fluctuating environmental conditions is given by:

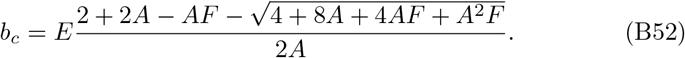

In the absence of temporal fluctuations in the environmental conditions (i.e., when *F* = 0) equation (B52) reduces to

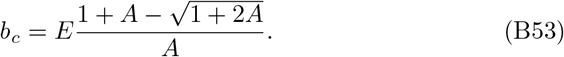

Recall that in habitats with a spatially constant environmental gradient, a population without the capacity for plasticity faces global extinction above the critical genetic gradient given by equation (B52) (or equation (B53) in static environmental conditions), whereas it successfully expands and adapts to the entire habitat below the critical genetic gradient. Conversely, in habitats with a spatially steepening environmental gradient, the critical genetic gradient (equations (B52)-(B53)) indicates the spatial position where, in the absence of plasticity, adaptation fails and range expansion stops.

Next, we turn to the model in which the expanding population has capacity for plasticity (see model details in Appendix B1). For this model, we first seek a gradient below which plasticity of zero is optimal (hereafter called the *critical plasticity gradient*). Recall from equation (B44) that zero plasticity is optimal at local environmental gradients *b*(*x*) such that:

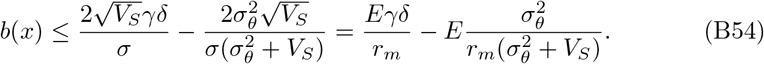

Note that the RHS of equation (B54) scales linearly with *γδ*, that is, it increases linearly when increasing the cost of plasticity. Conversely, a positive value of plasticity is optimal in positions where the environmental gradient *b*(*x*) is larger than the RHS of equation (B54). Thus, for a given cost of plasticity, there exists a minimal gradient, i.e., the critical plasticity gradient 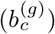 above which the optimal plasticity is positive

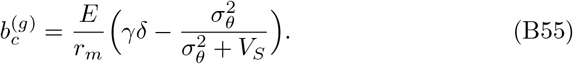

Recall that for a population without the capacity for plastic response in the adaptive trait, a range margin forms when the environmental gradient is equal to *b*_*c*_ given by equation (B52). For a population that has the capacity for plasticity, we have shown here that there is a critical plasticity gradient below which zero plasticity is optimal. When the critical plasticity gradient is larger than *b*_*c*_, plasticity in equilibrium is 0 at *b*_*c*_, and at all shallower gradients. In this case, thus, the population evolves as if it does not have the capacity for a plastic response in the adaptive trait. This is because, despite the fact that the population has the capacity for plasticity, plasticity would not improve its fitness, and hence it does not evolve (or, if it evolves, it does so transiently and it is not maintained). In other words, when the critical plasticity gradient 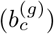 is steeper than the critical genetic gradient *b*_*c*_, adaptation of the population fails when *b*(*x*) = *b*_*c*_, and the range of the population corresponds to that determined in [11]. Otherwise, when the critical plasticity gradient 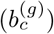 is shallower than the critical genetic gradient (*b*_*c*_), the range of the population with the capacity for plasticity is larger than when plasticity is absent.

Note that the cost of plasticity in equation (B54) determines the value of the critical plasticity gradient. This implies that there is a critical cost of plasticity *δ*_*c*_ for which the critical plasticity gradient equals the critical genetic gradient. To find this critical cost of plasticity (*δ*_*c*_), we require that equation (B55) evaluated at *δ*_*c*_ is equal to *b*_*c*_ (given by equation (B52)). Solving for *δ*_*c*_ yields

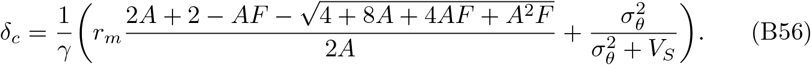

In conclusion, we find that, when *δ* > *δ*_*c*_, the population range-expansion dynamics are within *R*_0_ regime (as per the notations used in the main text), i.e., the regime where the capacity to (potentially) evolve plasticity does not increase the equilibrium range in comparison to when the capacity for plasticity is absent.

By contrast, when *δ* < *δ*_*c*_, the equilibrium range may be larger when a population has the capacity for plasticity, compared to a population that does not. In this case, there are two possibilities for the equilibrium range of the population when plasticity may evolve: a finite but larger range compared to the case without the capacity for plasticity (regime *R*_1_ in the main text), or an infinitely large range (accounted for by regime *R*_2_, as explained next; and see main text).

A necessary but not sufficient condition for a parameter combination to allow infinite range expansion is that the population can maintain a positive growth rate with a plasticity of 1, that is, when *r*_*m*_ > −*γ* ln(1 − *δ*) (equation (B15)). If this inequality is not met, then there must be an upper limit to plasticity that may evolve. As a consequence, for a large enough steepness of the environmental gradient, adaptation must fail. In other words, when *δ* < *δ*_*c*_ but *r*_*m*_ ≤ −*γ* ln(1 − *δ*), the parameters are in the *R*_1_ regime. Conversely, when *δ* < *δ*_*c*_ and *r*_*m*_ > −*γ* ln(1 − *δ*), the parameters are in the *R*_2_ regime. For parameters within regime *R*_2_ the population may have an infinite range when the population attains the optimal plasticity of 1 in equilibrium, which may not always be the case (recall that we only found a necessary but not sufficient condition for this to be true). For example, when the region of the environmental gradient where intermediate values of plasticity are optimal is narrow and when dispersal is sufficiently strong relative to selection on plasticity, the allele frequencies for plasticity may change slower across space than the optimal plasticity (using the arguments in [85] *albeit* in a model without plasticity). With this caution in mind, we use *r*_*m*_ = −*γ* ln(1 − *δ*) to separate the *R*_1_ regime from the *R*_2_ regime, where infinite range expansion is possible either in this entire region, or in a sub-region. Clearly, infinite range expansion is expected in the absence of any plasticity costs. This is because, in this case, all individuals may have plasticity of 1 without any penalty, and with this attain perfect adaptation everywhere (assuming that the non-plastic component of the phenotype remains, on average, the same as in the expansion source, i.e., 0).

We emphasise that the calculations in this appendix are based on a number of simplifying assumptions. Among them is the assumption that plasticity is constant over time and locally in space (and that it is the same for all local individuals). When plasticity varies between individuals within a local population, the optimal population mean of plasticity may be different from the optimal plasticity found above (as also found in our simulations, e.g. figures 3 C, C14). In addition, a potential covariance between plasticity and the non-plastic component of genetic adaptation may alter the optimal population mean of plasticity but we neglected this here. Furthermore, as discussed above, dispersal between neighbouring demes may alter the local population mean of plasticity. In conclusion, the local population mean of plasticity that a population attains in the long run may not be equal to the optimal mean population plasticity approximated here. But, despite this, our simulation and analytical results agree relatively well (this is further discussed in the main text).

Importantly, when the cost of plasticity is above the critical cost, given by equation (B56), the optimal plasticity is zero for all environmental gradients up to the critical genetic gradient as defined in [11]. As a consequence, dispersal does not alter plasticity in equilibrium, because it is expected to be the same (i.e., zero) in all demes. Furthermore, there is negligible covariance between plasticity and the non-plastic component of genetic adaptation, because the variance in plasticity is expected to be low. In this case, our analysis shows that the model outlined in the main text reduces to the model in [11] (see also [40]). Thus, equation (B56) gives an approximate condition for when the capacity for plasticity cannot make the equilibrium range of the population larger than the range of a population without the capacity for plasticity.

The results derived here guided our choice of the parameter values examined in the individual-based model presented in the main text. Furthermore, they aided the interpretation and qualitative understanding of our simulation results. The results from the individual-based model and their interpretation are discussed in detail in the main text.

## Appendix C Additional simulation results

In this appendix, we present additional simulation results that are relevant for the interpretation of our findings.

## C.1 Range expansion without plasticity

Figure C1 shows the population size 100, 000 generations after the start of range expansion for populations without the capacity for plasticity. When there were no temporal fluctuations in the optimal phenotype, we obtained the expected results for range expansion without plasticity along steepening environmental gradients (figure C1 A) [40, 11, 41]. When the phenotypic optimum fluctuated in time, the population size was reduced in comparison to when the phenotypic optimum was static (figures C1 B-D) in agreement with equation (B49) and [14]. As a consequence of the reduced population size, the equilibrium range was reduced in comparison to the expectation in a temporally static environment (purple crosses in figure C1).

## C.2 Static environment

Here, we present the simulation results for range expansions where plasticity was allowed to evolve and the environmental conditions were temporally static. Figures C2-C4 show simulation results obtained at the end of the burn-in period for temporally static environments. In this case, plasticity was nearly uniformly zero for all costs of plasticity examined (figure C2). The cline patterns for the loci coding for the non-plastic component of the phenotype were similar to the expected clines in the absence of plasticity (compare black and red lines in figure C3). However, there was one locus that was almost fully heterozygous in the centre of the habitat for all parameters (figure C3), even though the environmental gradient was zero in the centre. This is likely due to the fact that the model had an odd number of loci, and thus a population that on average was perfectly adapted had to be heterozygous at one locus. The frequencies for alleles coding for plasticity did not seem to have any obvious pattern (figure C4).

Figure C5 shows the evolution of plasticity during range expansion for different values of the plasticity cost parameters *δ* and *γ*, and of the local carrying capacity, *K*. When the cost of plasticity was above the critical cost of plasticity (see Results in the main text) almost no plasticity evolved throughout the range (figures C5A-C), although low positive plasticity evolved in the range margin for cost parameters close to the critical cost (figure C5 C). By contrast, when the cost of plasticity was below the critical cost of plasticity, high plasticity evolved during range expansion (figure C5 D). In this case, positive plasticity initially evolved in the range margin, and thereafter plasticity started increasing towards the centre of the range.

Figure C6 shows the population size and plasticity attained 100, 000 generations after the start of range expansion. The parameters in figure C6 correspond to those in figure C5. When the cost of plasticity was above the critical cost of plasticity, very little plasticity evolved, although a slightly increased plasticity occurred near the range margins (red line in figures C6 A-C). In all cases, the equilibrium population size agreed with the prediction in [11] (blue lines in C6 A-C). Conversely, when the cost of plasticity was below the critical cost, high plasticity evolved towards the range margins (red line in figure C6 D). As a consequence, the population size reached a plateau (of about 60 individuals) approximately 50 demes before the edge of the habitat (blue line in figure C6 D; whereas, in the absence of plasticity, the population size would sharply decay towards zero approximately 50 demes before the edge of the habitat).

**Figure C1:**
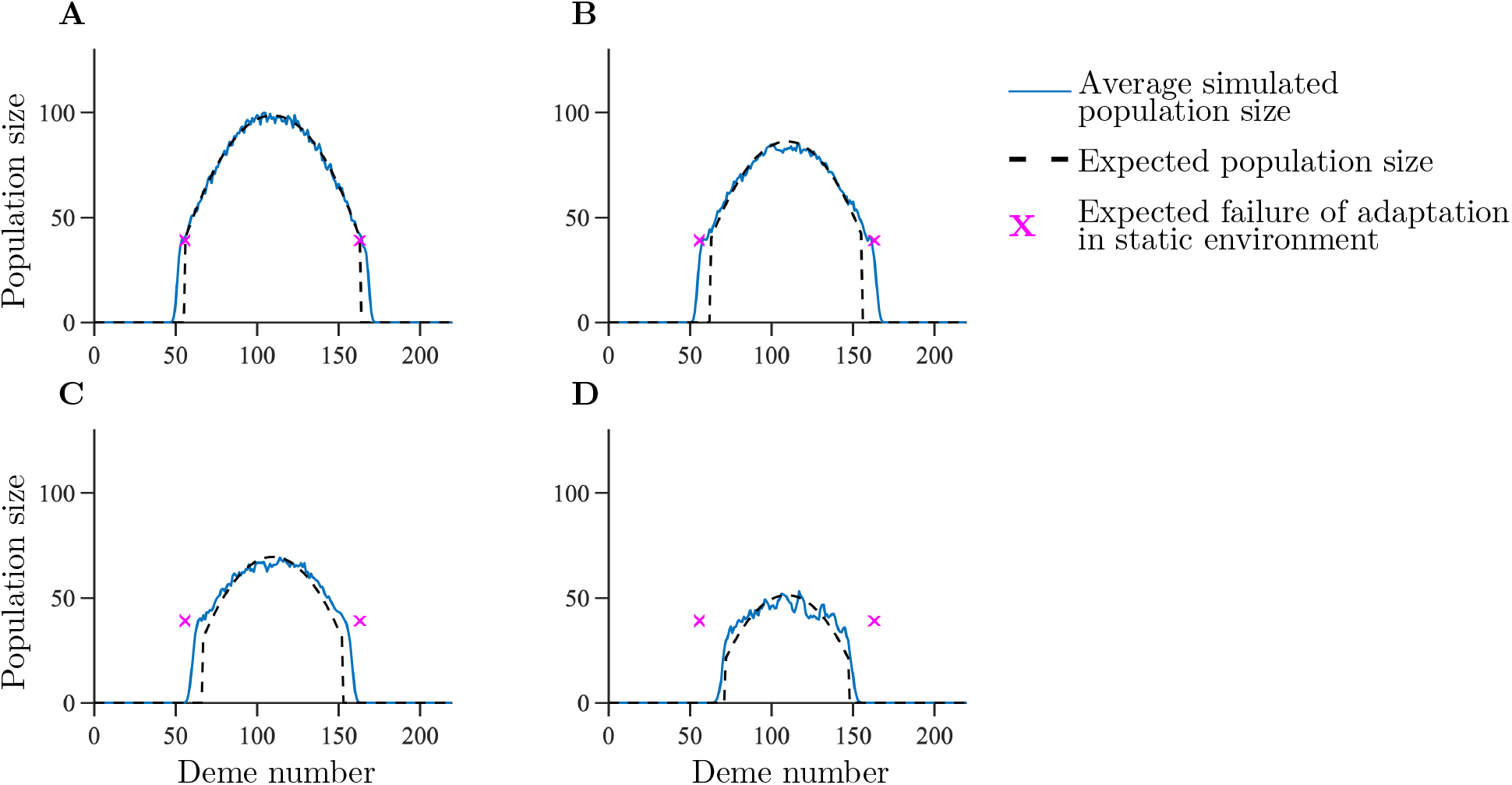
Average population size attained 100, 000 generations after the start of range expansion for a population without the capacity for plasticity. The panels differ by the parameter *σ*_*θ*_: *σ*_*θ*_ = 0 (A), 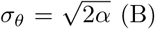 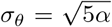 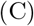, 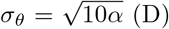. The blue line shows the realised population size 100,000 generations after the start of range expansion, averaged over 100 realisations. The black dashed line shows the population size given by equation (B49). The purple crosses indicate the expected failure of adaptation when *σ*_*θ*_ = 0. Remaining parameters: *K* = 100, *r*_*m*_ = 1, *V*_*S*_ = 2, *μ* = 10^−6^, *σ* = 1.

In figure C7, the evolution of plasticity, as well as the population size and plasticity attained 200, 000 generations after the start of range expansion is shown for *γ* = 0.25. The remaining parameters are the same as in figure 2 B in the main text. Figure C7 shows that, for smaller *γ* (i.e., more concave cost-related function), the range may be much larger than in the absence of plasticity, even though the range is finite.

Figure C8 shows an example of the spatial pattern of allele frequencies for the alleles coding for the non-plastic component of the phenotype 100, 000 generations after the start of range expansion. The spatial pattern of allele frequencies for the alleles coding for the non-plastic component of the phenotype corresponded to a series of staggered clines. These had the same width as the clines for a population without the capacity for plasticity (compare the black lines, corresponding to the simulation results, to the theoretical expectation shown in red lines in figure C8 A). The spacing between the clines was, however, different to the expected spacing for a population without plasticity. As for a population without the capacity for plasticity, the clines were sparse in the centre of the habitat due to the shallow environmental gradient there. However, the clines were also sparser in regions where plasticity was the main mechanism for adaptation (compare the region between demes 10 and 50 in panel A to the same region in panel B).

When high plasticity evolved, the spatial pattern of allele frequencies for alleles coding for plasticity formed cline-like patterns that were increasing from the centre towards the edges on both sides, rather than increasing from one to another edge (figure C8 B).

## C.3 Environment with optimum that fluctuates in time

Here, we present the simulation results for range expansions where plasticity was allowed to evolve and the environmental conditions were fluctuating in time.

Figures C9-C11 show simulation results obtained at the end of the burn-in period for temporally fluctuating environments. With temporal fluctuations in the optimal phenotype, no (or very low) plasticity evolved when the cost of plasticity was above the critical cost (figures C9 A, C9 B, and C9 D). However, plasticity evolved during the burn-in period when the cost of plasticity was below the critical cost (C9 C, C9 E, F, G, H and C9 I). Note that plasticity was approximately equally strong in all demes throughout the habitat (the black lines in figure C9 are approximately straight and parallel to the *x*-axis). In some cases (e.g., figures C9 C, C9 E, and C9 G), plasticity that evolved in the simulations was slightly higher than expected from equation (B41). This is possibly because equation (B41) relies on the assumption of a spatially homogeneous environment, whereas the habitat contains a (shallow) environmental gradient during the burn-in period. In these cases, the joint effect of spatial and temporal variability may cause the optimal plasticity to be positive (rather than zero, as expected from temporal fluctuations alone). The spatial patterns of allele frequencies for the alleles coding for the non-plastic component of the phenotype were more noisy than for static environments (figure C10; compare to figure C3). As for when the environmental conditions were static, there was no evident spatial pattern in the allele frequencies for plasticity (figure C11).

Figure C12 shows the evolution of plasticity during range expansion in the presence of temporal fluctuations of the optimal phenotype. As expected from equation (B49), when the cost of plasticity was above the critical cost, the range was smaller than the equilibrium range in temporally static environments (figures C12 A, C12 C, and C12 E). By contrast, when the cost of plasticity was below the critical cost, temporal fluctuations, instead, increased the range ((and range expansion was faster when temporal fluctuations were larger; compare panels B, D, and F in figure C12).

Figure C13 shows the population size and plasticity attained 100, 000 generations after the start of range expansion. The parameters in figure C13 correspond to those in figure C12. When the cost of plasticity was above the critical cost, almost no plasticity had evolved 100, 000 generations after the start of range expansion (red lines in figures C13 A, C13 C, and C13 E). In this case, the population size and the range were decreased, in comparison to the expected population size and range for temporally static environmental conditions (blue line in figures C13 A, C13 C and C13). By contrast, high plasticity evolved when the cost of plasticity was below the critical cost (red lines in figures C13 B, C13 D, and C13). In this case, the population size was almost constant throughout the habitat as expected due to the high plasticity (blue lines in figures C13 B, C13 D, and C13 F).

## C.4 Additional simulations

In figure C14, the population size and plasticity attained 200, 000 generations after the start of range expansion are shown in a habitat with a phenotypic optimum that changes linearly in space. The gradient is such that the optimal plasticity according to equation (B25) is equal to 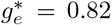. Notably, there is a shallow gradient in plasticity 200, 000 generations after the start of range expansion. This gradient is likely caused, in part, due to stronger selection on plasticity along an environmental gradient (cf. [23]), together with edge effects caused by the finite number of loci used in the simulations. Although the gradient in plasticity may be a transient effect and plasticity may level out in a longer run (as argued in [25]), the results in figure C14 indicate that the fitness benefit to further decrease plasticity in the edges or increase it in the centre is too weak to reach the optimal plasticity within 200, 000 generations.

In figure C15, the evolution of plasticity and the population size and plasticity attained 100, 000 generations after the start of range expansion is shown for the same parameter values as in figure 2 B, except that here the number of loci underlying plasticity was smaller, and the magnitude of the allele effect sizes at these loci was proportionately larger (details in Methods in the main text). In this case, much higher plasticity evolves at the range margin (compare figure C15 to figure 2 B). This is because, due to larger effect sizes at loci underlying plasticity, selection per locus is stronger in the case shown in figure C15 than in figure 2. This favours the evolution of higher plasticity in the range margin, where the population experiences continued directional selection to restore the mean phenotype to the local optimum [9]. However, despite stronger selection for plasticity, there is a limit to the amount of plasticity that can evolve and the range expansion dynamics fall within regime *R*_1_, as our analytical analysis shows.

These and other results we obtained are further discussed in the main text.

**Figure C2:**
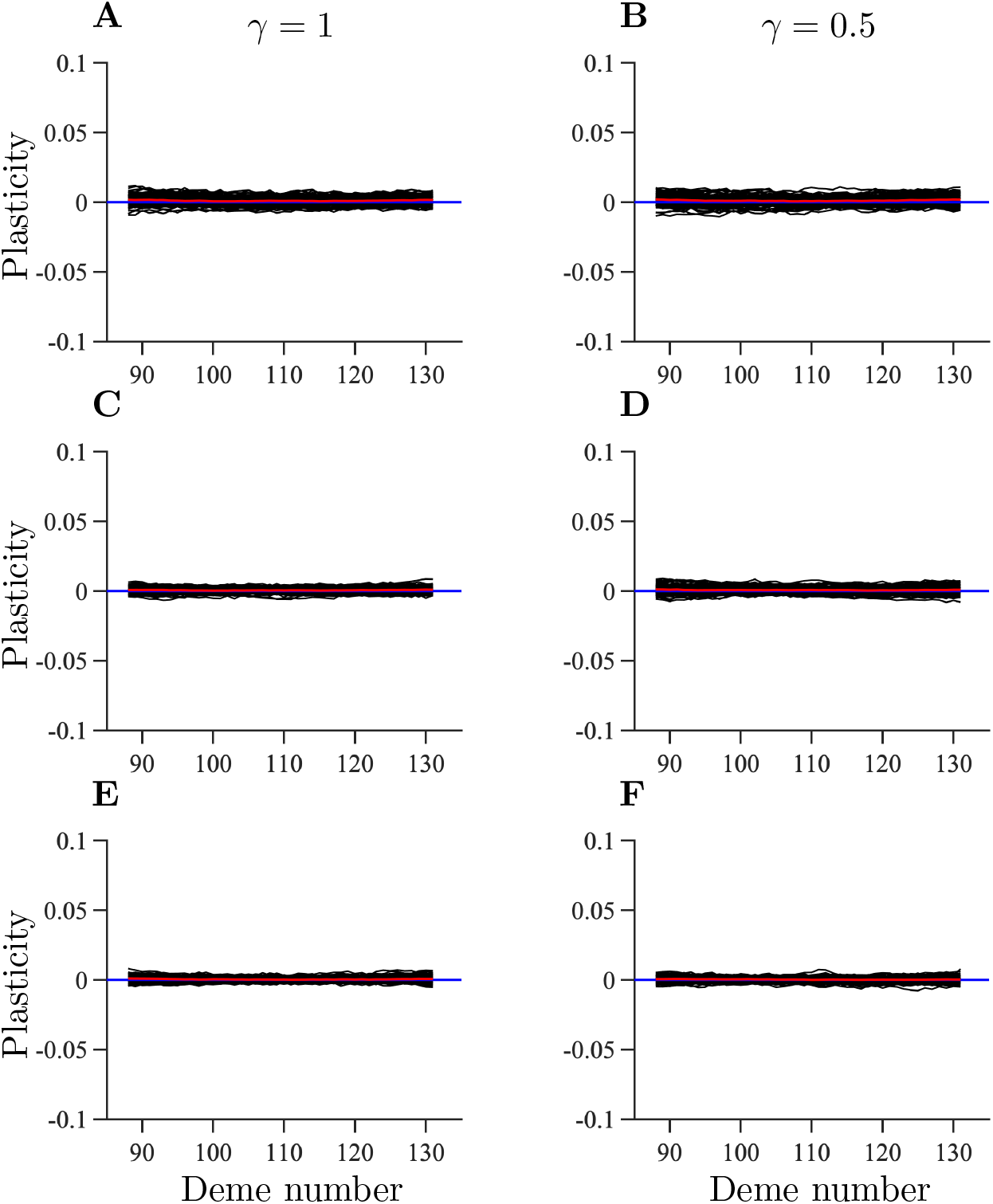
Average plasticity in static environments at the end of the burn-in period. The panels differ by the parameters *γ* and *δ*: *γ* = 1, *δ* = 0.25 (A), *γ* = 0.5, *δ* = 0.5 (B), *γ* = 1, *δ* = 0.6 (C), *γ* = 0.5, *δ* = 0.9 (D), *γ* = 1, *δ* = 0.75 (E), and *γ* = 0.5, *δ* = 1.35 (F). The black lines show the population-average plasticity for individual realisations. The red line shows the total average over 100 realisations. The blue line shows the analytically calculated plasticity in equilibrium in an environment that is spatially homogeneous. Remaining parameters: *K* = 100, *r*_*m*_ = 1, *V*_*S*_ = 2, *μ* = 10^−6^, *σ* = 1, *L* = 799, *α* = 0.3162, *β* = 0.0013, and *σ*_*θ*_ = 0. In each case 100 realisations were performed.

**Figure C3:**
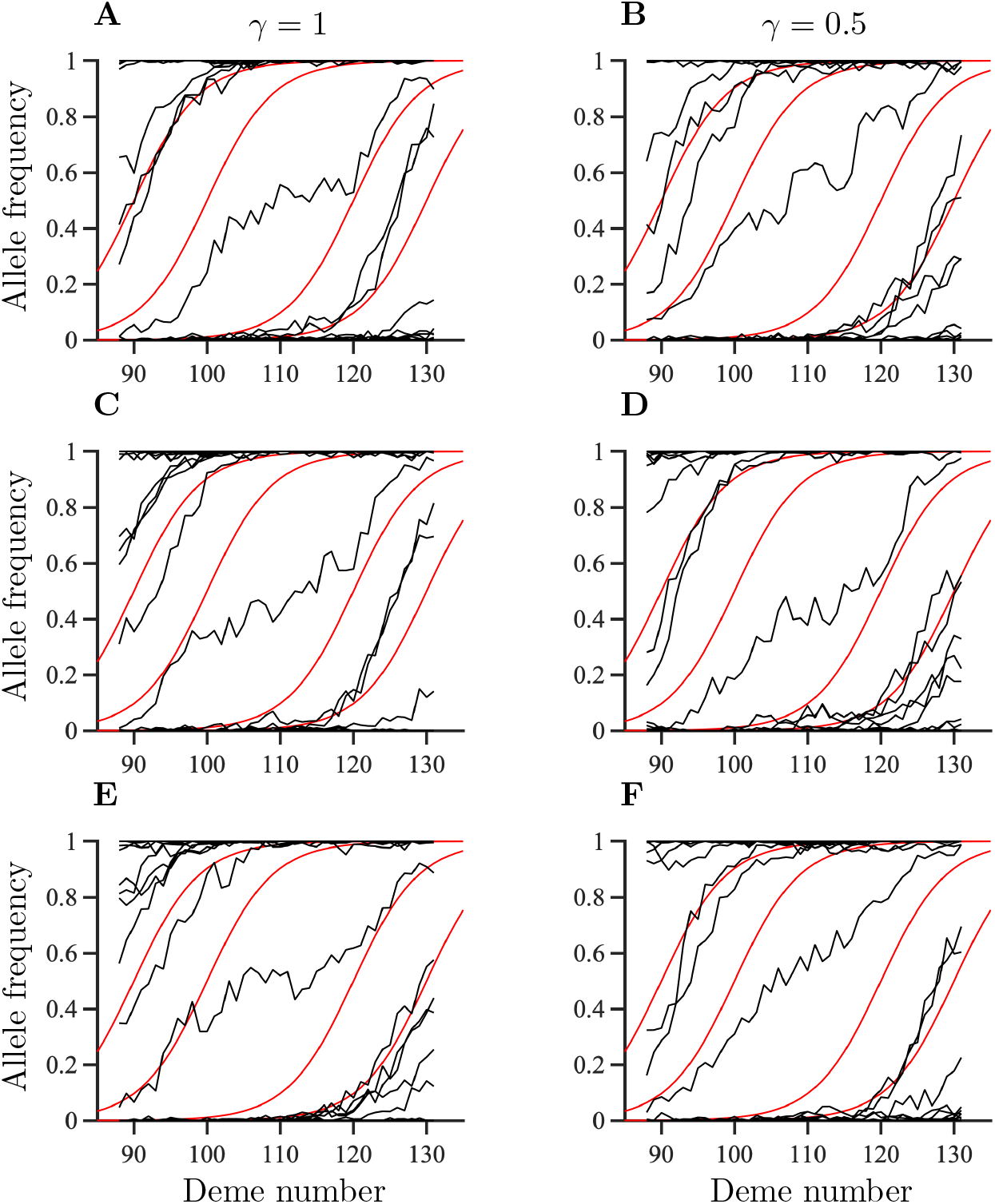
Spatial patterns of allele frequencies for the non-plastic component of the phenotype in static environments at the end of the burn-in period for single randomly chosen realisations. The parameters shown in panel A-F correspond to those in figure C2. The black lines show the realised allele frequencies for the alleles coding for the non-plastic component of the phenotype. The red lines show illustrative examples of theoretically expected clines in allele frequencies: *p*_*z,j*_ = 1/(1 + exp(−4(*x* − *c*_*j*_)/*w*)), where 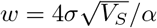 and *x* and *c*_*j*_ denote the spatial position and the centre of the cline, respectively. Here, the centres of the clines are located in demes 90, 100, 120, and 130.

**Figure C4:**
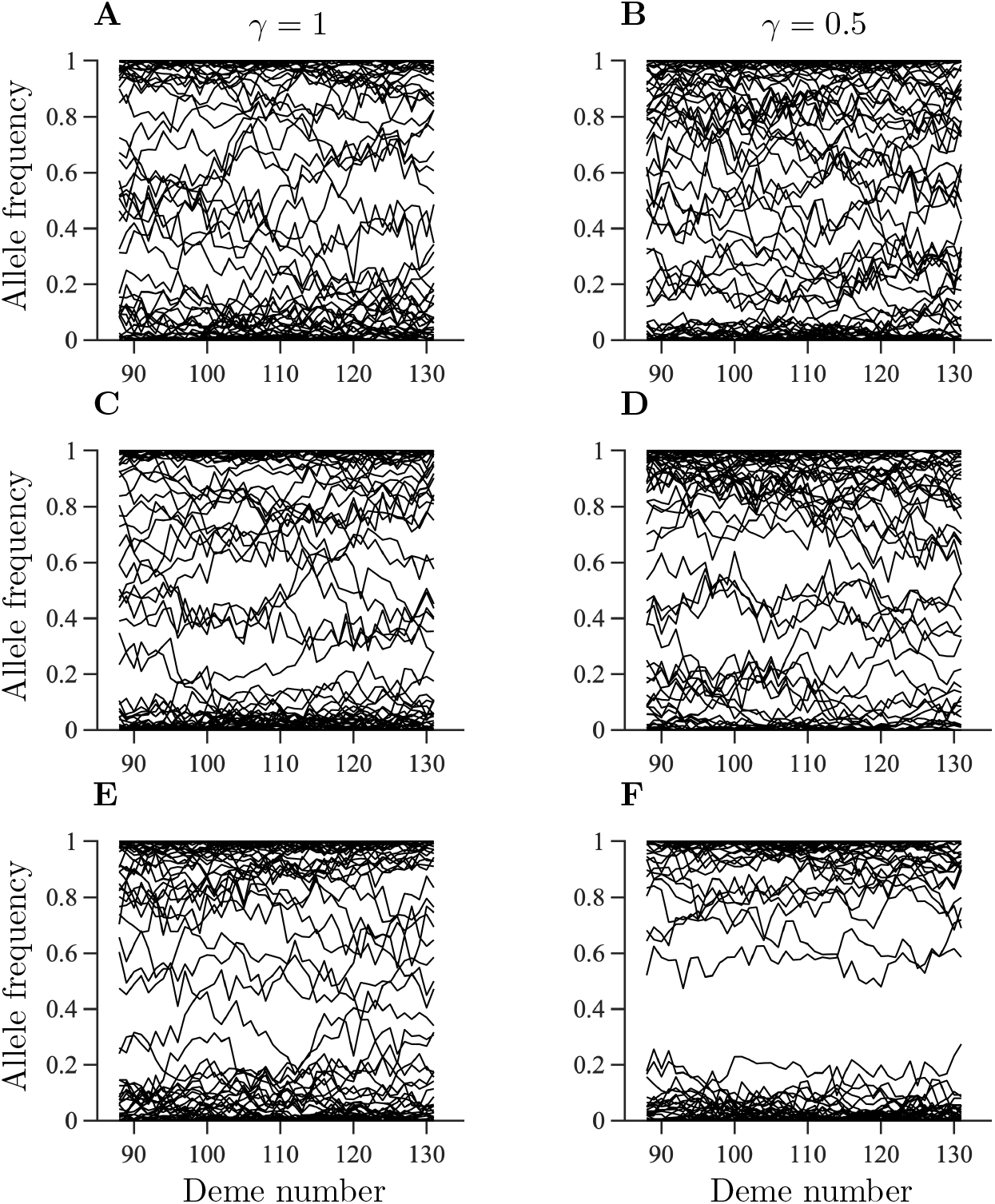
Spatial patterns of allele frequencies for the alleles coding for plasticity in static environments at the end of the burn-in period for single randomly chosen realisations. The parameters shown in panel A-F correspond to those in figure C2.

**Figure C5:**
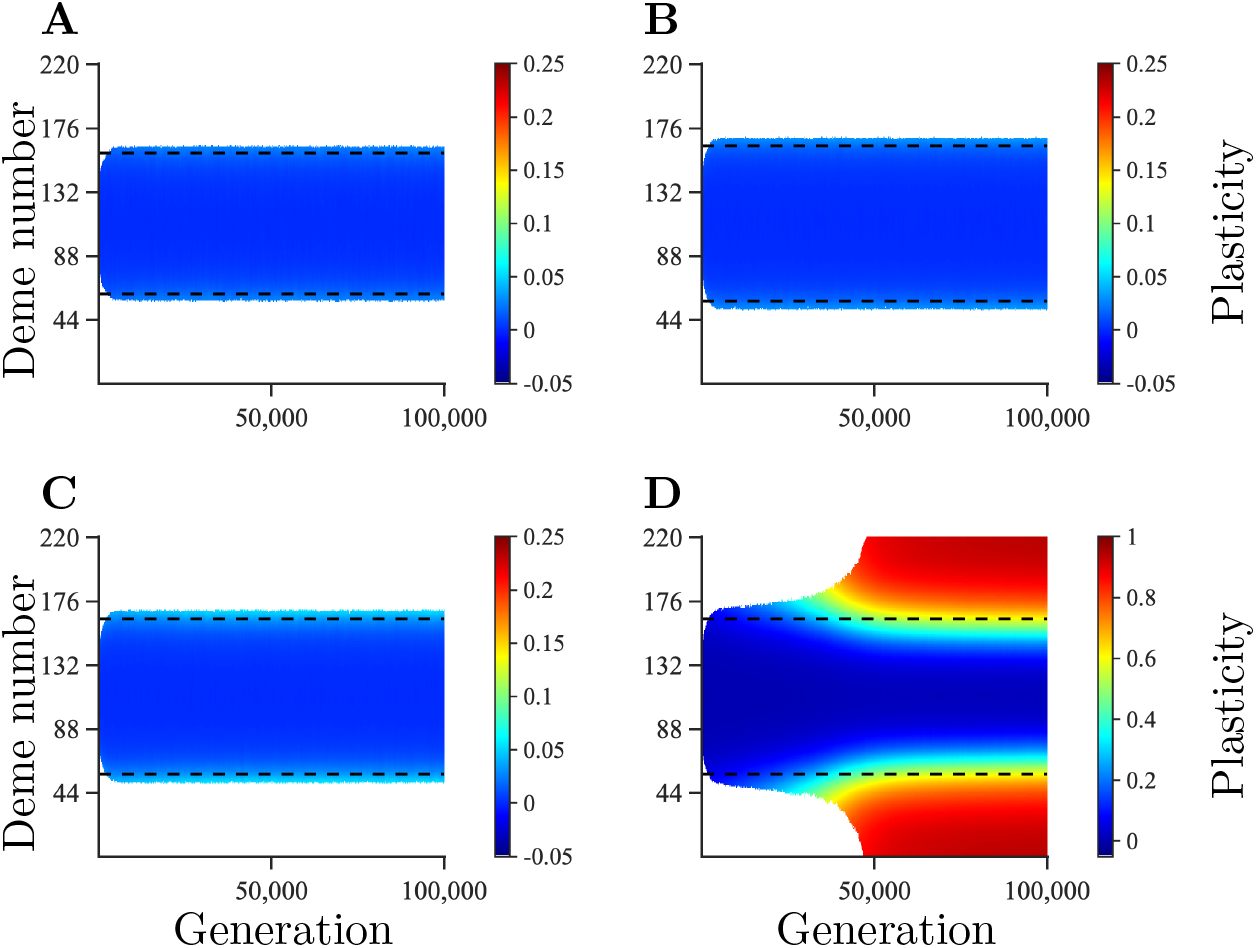
Temporal and spatial evolution of plasticity averaged over 100 realisations during range expansion in a habitat with temporally static environmental conditions. The dashed lines denote where adaptation is expected to fail for a population without plasticity. The panels differ by the parameter *δ* and *K*: *δ* = 0.5, *K* = 50 (A), *δ* = 0.75, *K* = 100 (B), *δ* = 0.6, *K* = 100 (C), and *δ* = 0.25, *K* = 100 (D). Remaining parameters: *r*_*m*_ = 1, *V*_*S*_ = 2, *μ* = 10^−6^, *σ* = 1, *L* = 799, *α* = 0.3162, *β* = 0.0013, *γ* = 1, and *σ*_*θ*_ = 0.

**Figure C6:**
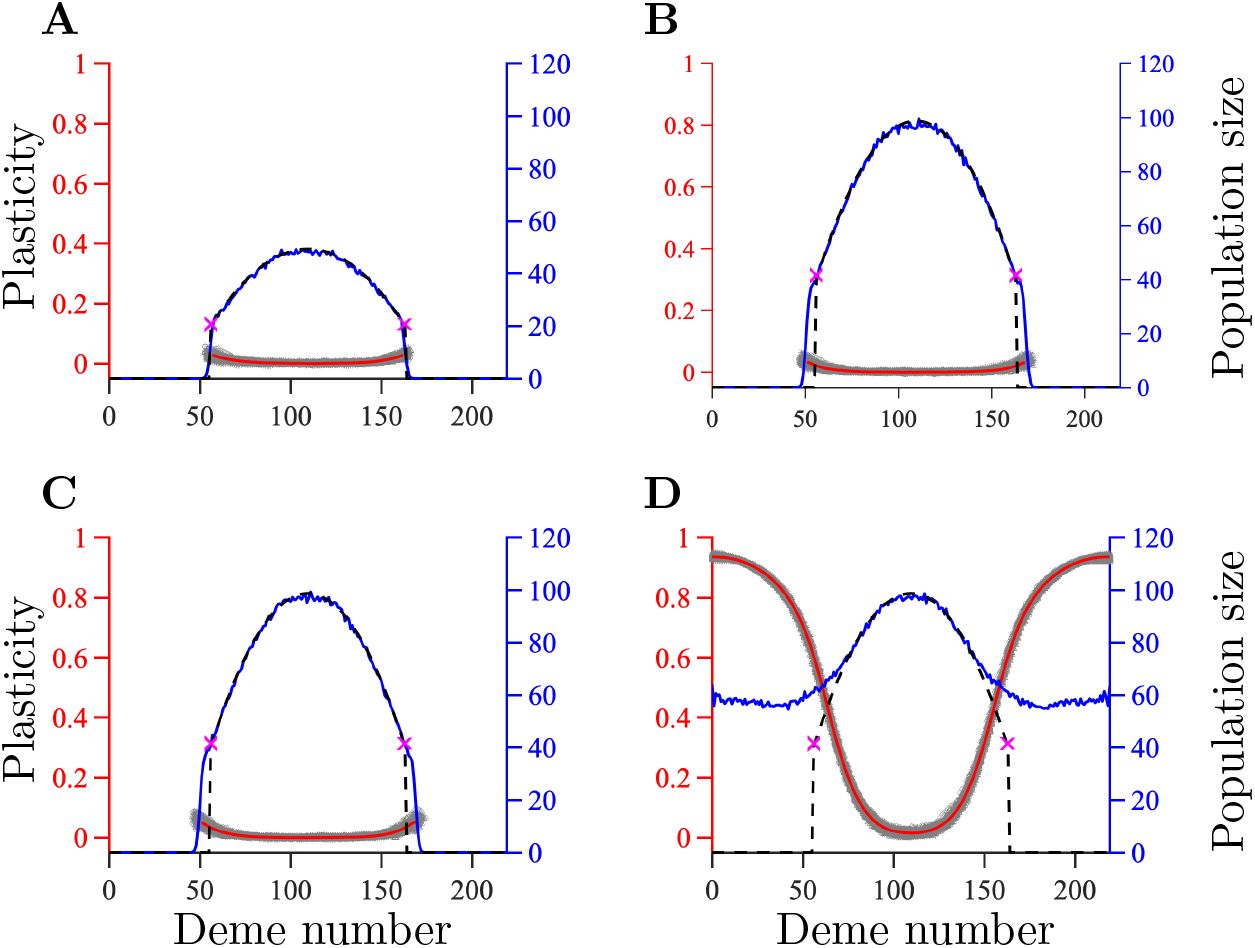
Population size and plasticity 100, 000 generations after the start of range expansion in a habitat with temporally static environmental conditions. The results in panels A-D corresponds to those in panels A-D in figure C5, respectively. The red axis and red line show plasticity averaged over 100 realisations, the grey area indicates the spread of plasticity values obtained in different realisations. The blue axis and blue line show the population size, averaged over 100 realisations. The expected population size, and the deme where adaptation is expected to fail in the absence of plasticity are shown by the dashed line, and purple crosses, respectively.

**Figure C7:**
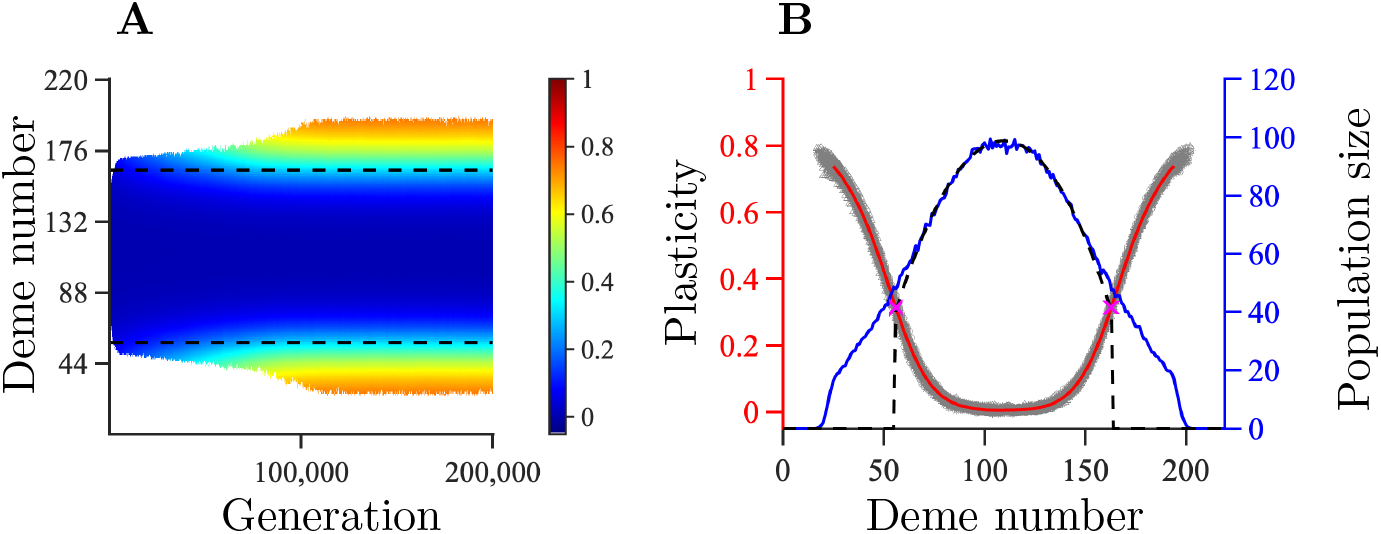
Range expansion in a habitat with temporally static environmental conditions and shape parameter *γ* = 0.25 for the function related to the cost of plasticity. Temporal and spatial evolution of plasticity averaged over 100 realisations during range expansion (A). Population size and plasticity 200, 000 generations after the start of range expansion (B). The dashed lines in panel A denote where adaptation is expected to fail for a population without plasticity. The red axis and red line in panel B show plasticity averaged over 100 realisations, the grey area indicates the spread of plasticity values obtained in different realisations. The blue axis and blue line show the population size, averaged over 100 realisations. The expected population size, and the expected failure of adaptation in the absence of plasticity are shown by the dashed line, and purple crosses, respectively. Remaining parameters: *K* = 100, *r*_*m*_ = 1, *V*_*S*_ = 2, *μ* = 10^−6^, *σ* = 1, *L* = 799, *α* = 0.3162, *β* = 0.0013, *δ* = 1.1, and *σ*_*θ*_ = 0.

**Figure C8:**
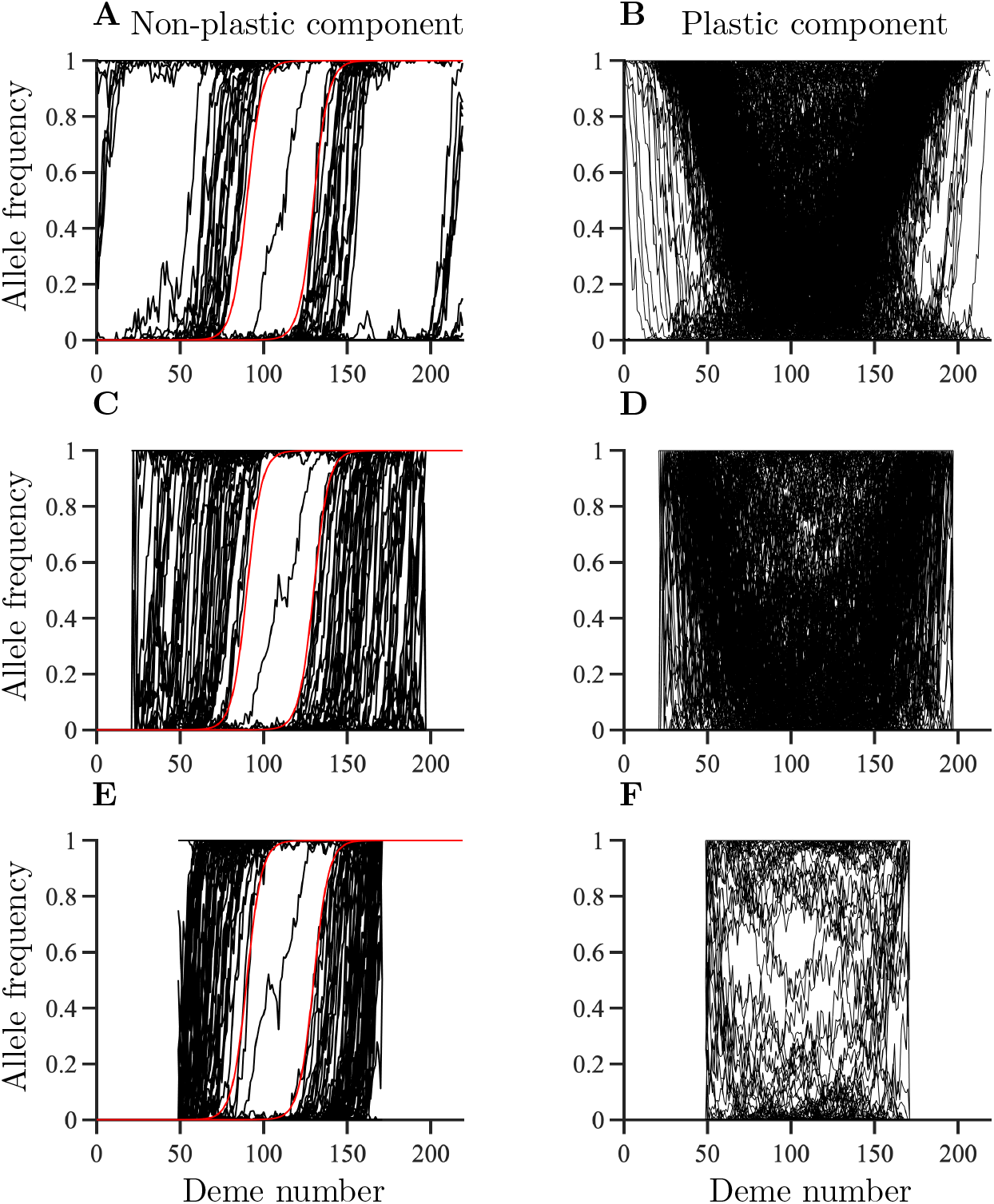
Examples of spatial patterns of allele frequencies for alleles coding for the non-plastic component of the phenotype (A, C, and E) and plasticity (B, D, and F). The parameters in panel A-B correspond to those in figures C5 D and C6 D (cost parameters *γ* = 1, *δ* = 0.25). The parameter values in panel C-D correspond to those in figure C7 (cost parameters *γ* = 0.25, *δ* = 1.1). The parameters in panel E-F correspond to those in figures C5 A and C6 A (cost parameters *γ* = 1, *δ* = 0.75). The allele frequencies shown in this figure were recorded 100, 000 generations after the start of range expansion for a single randomly chosen realisation. Black lines show realised allele frequencies. Two examples of analytical predictions for a cline, *p*_*z,j*_(*x*) = 1/(1+exp(−4(*x*−*c*_*j*_)*/w*)) (where *x* denotes spatial position, *c*_*j*_ denotes the centre of the cline, and 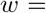 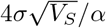, are shown in red in panel A, C and E. Remaining parameters: *K* = 100, *r*_*m*_ = 1, *V*_*S*_ = 2, *μ* = 10^−6^, *σ* = 1, *L* = 799, *α* = 0.3162, *β* = 0.0013, and *σ*_*θ*_ = 0.

**Figure C9:**
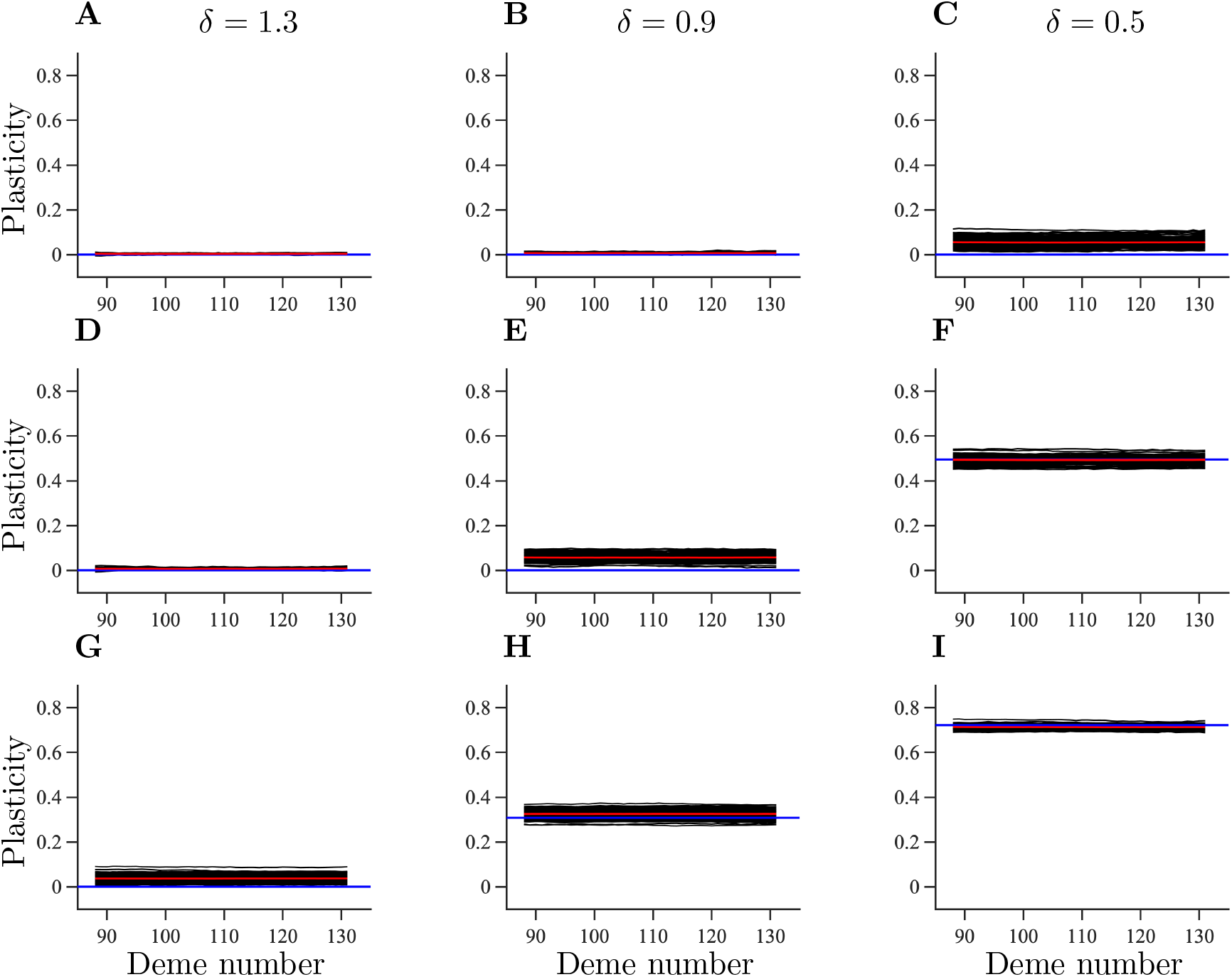
Plasticity at the end of the burn-in period in temporally fluctuating environments. The panels differ by the parameters *σ*_*θ*_ and *δ*: 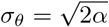, *δ* = 1.3 (A), 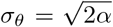, *δ* = 0.9 (B), 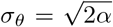, *δ* = 0.5 (C), 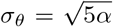, *δ* = 1.3 (D), 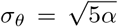, *δ* = 0.9 (E), 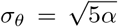, *δ* = 0.5 (F), 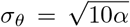, *δ* = 1.3 (G), 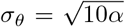, *δ* = 0.9 (H), and 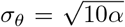, *δ* = 0.5 (I). Refer to the caption of figure C2 for an explanation of what the different lines represent. All panels in this figure have *γ* = 0.5. Remaining parameters are the same as in figure C2.

**Figure C10:**
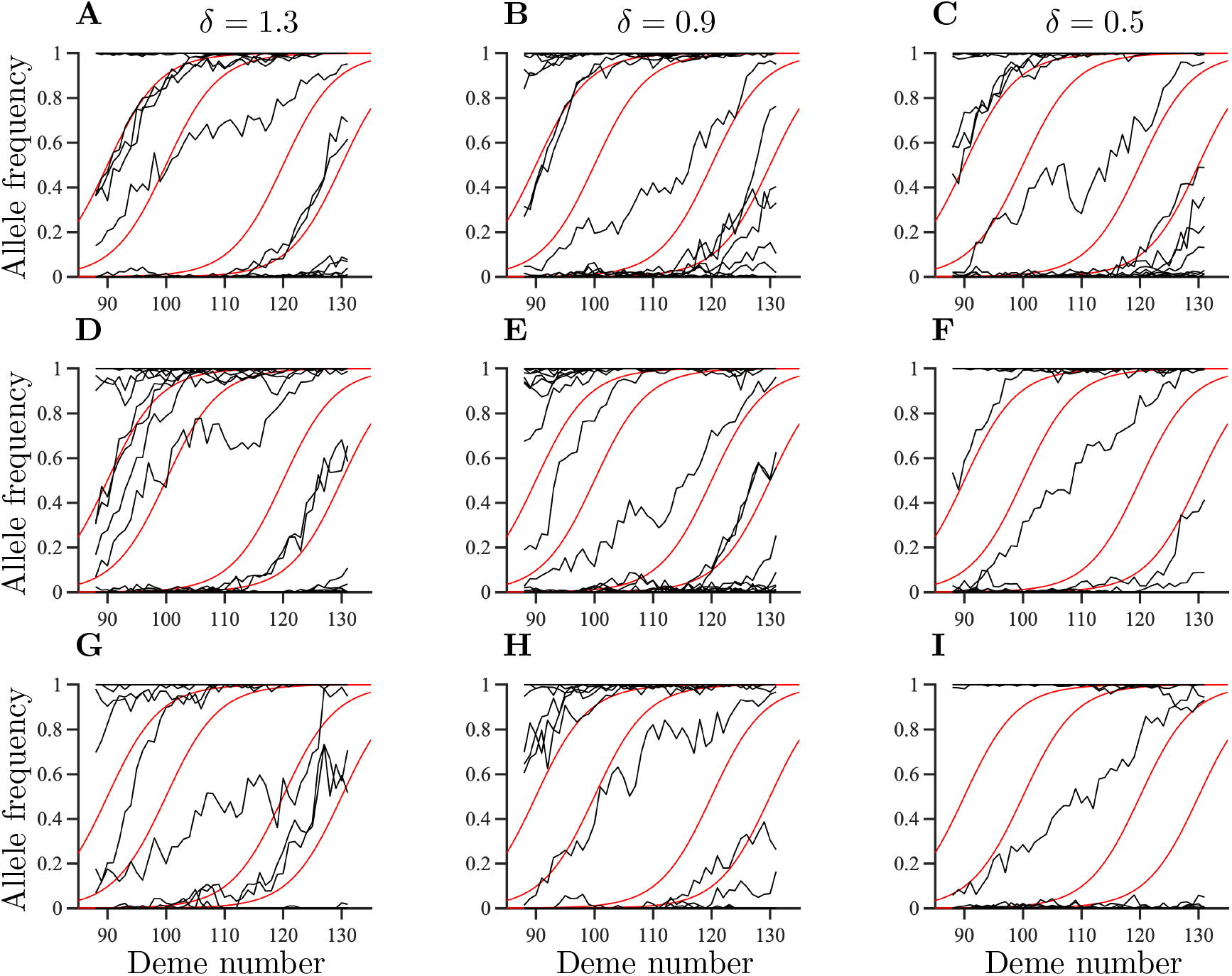
Spatial patterns of allele frequencies for the non-plastic component of the phenotype in temporally fluctuating environments at the end of the burnin period for single randomly chosen realisations. The parameters shown in panel A-I correspond to those in figure C9. Refer to the caption of figure C3 for an explanation of what the different lines represent.

**Figure C11:**
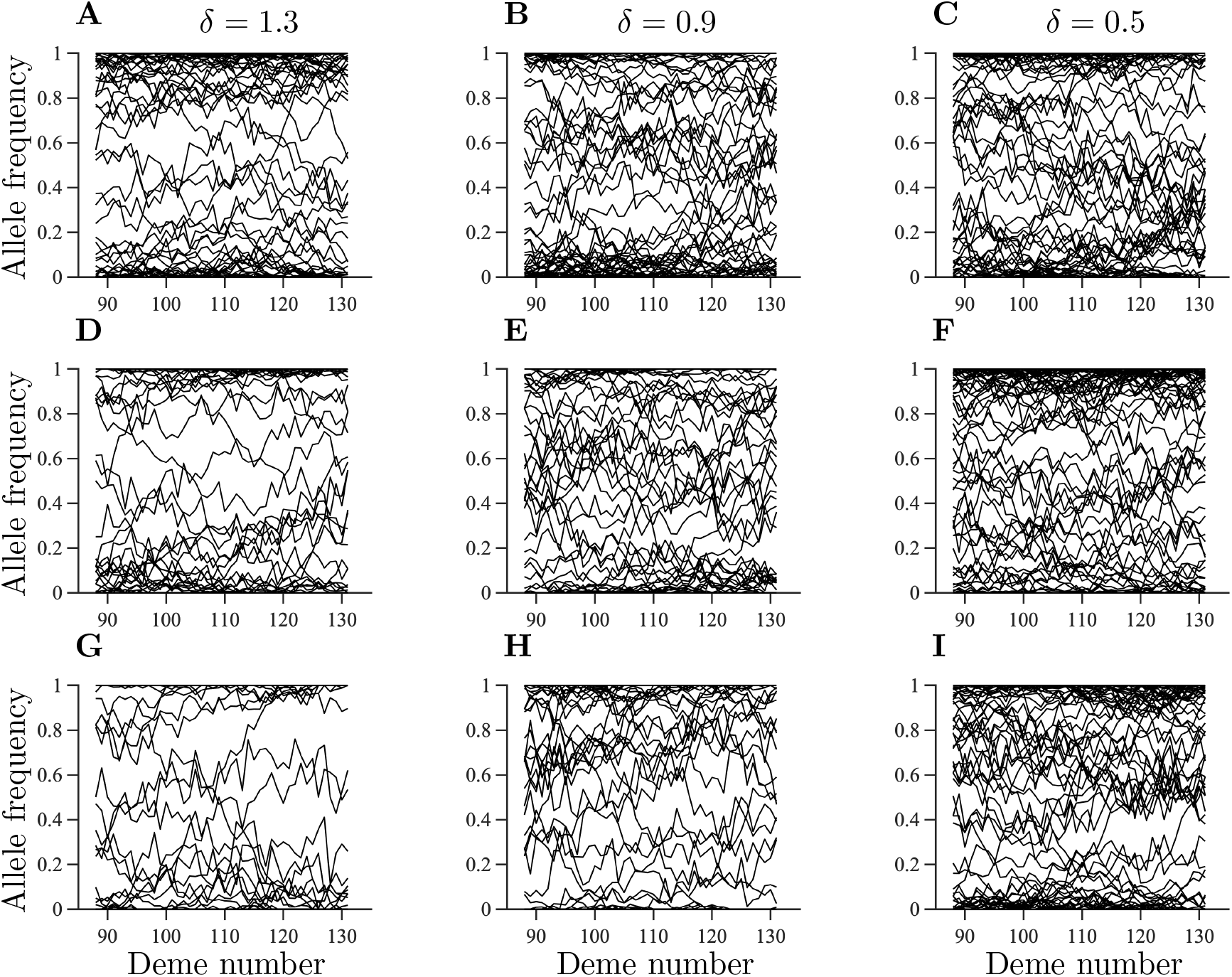
Spatial patterns of allele frequencies for the alleles coding for plasticity in temporally fluctuating environments at the end of the burn-in for single randomly chosen realisations. The parameters shown in panel A-I correspond to those in figure C9. Refer to the caption of figure C4 for an explanation of what the lines represent.

**Figure C12:**
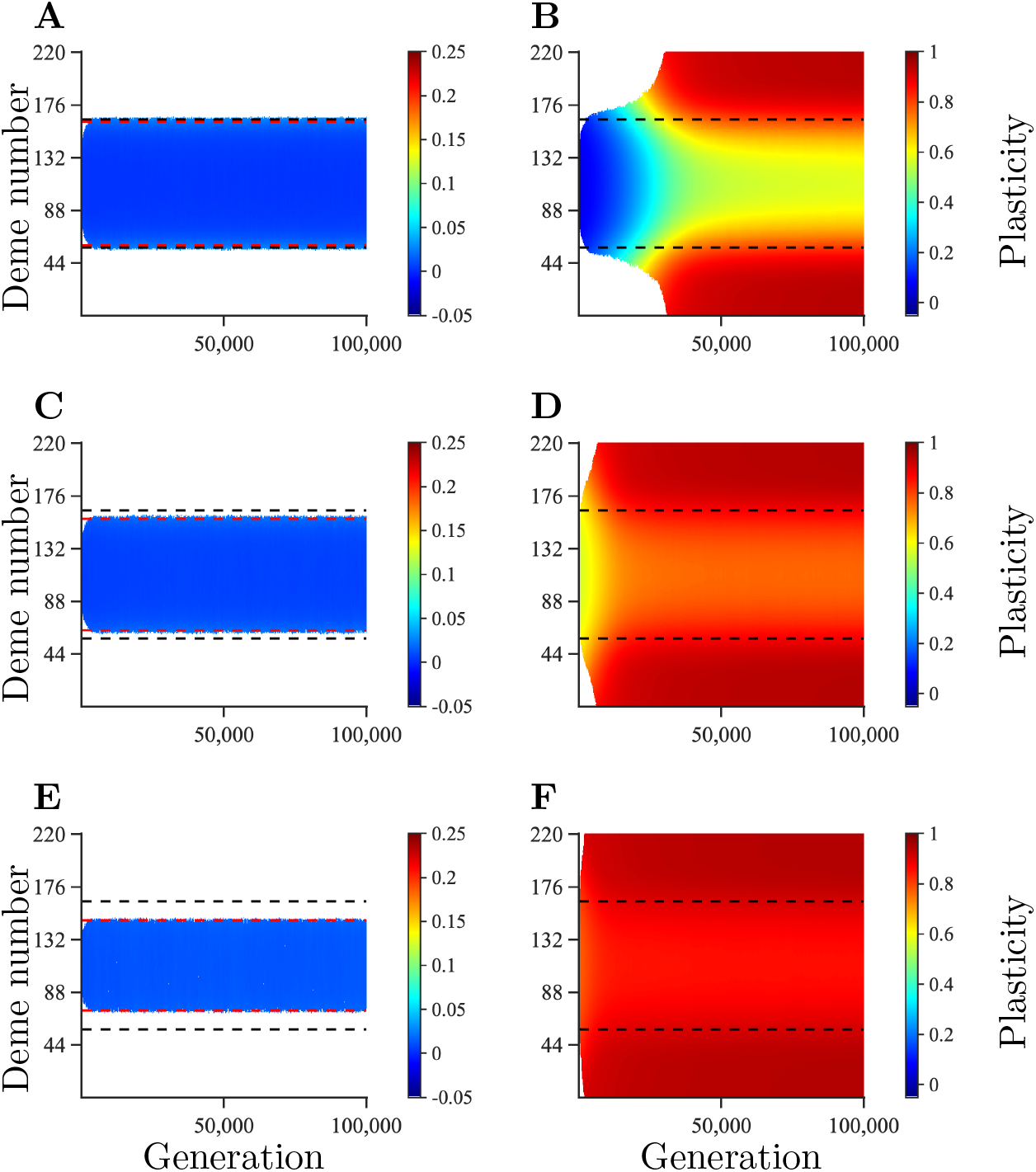
Temporal and spatial evolution of plasticity averaged over 100 realisations during range expansion in a habitat with temporally fluctuating environmental conditions. For comparison, the black dashed lines denote where adaptation is expected to fail for a population without plasticity in a temporally static environment. The red dashed lines show the expected failure of adaptation for a population without plasticity in a temporally fluctuating environment. The panels differ by the parameters *δ* and *σ*_*θ*_: *δ* = 0.75, 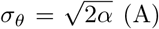, *δ* = 0.25, 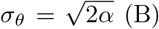, *δ* = 0.75, 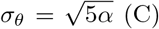, *δ* = 0.25, 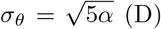, *δ* = 0.75, 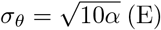, and *δ* = 0.25, 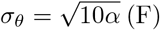. Remaining parameters: *K* = 100, *r*_*m*_ = 1, *V*_*S*_ = 2, *μ* = 10^−6^, *σ* = 1, *L* = 799, *α* = 0.3162, *β* = 0.0013, and *γ* = 1.

**Figure C13:**
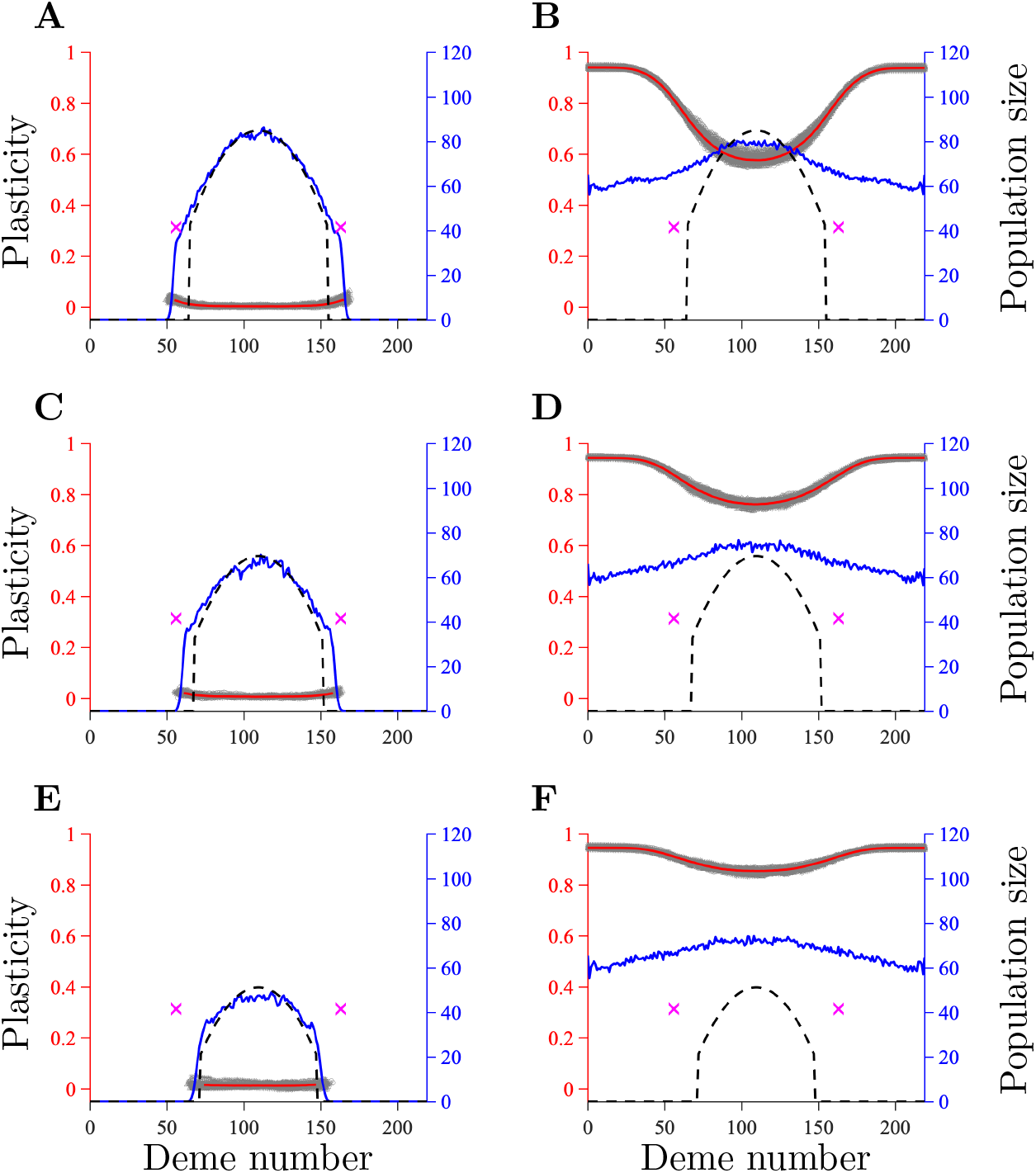
Population size and plasticity 100, 000 generations after the start of range expansion in a habitat with temporally fluctuating environmental conditions. The results in panels A-F correspond to those in panels A-F in figure C12, respectively. The expected population size for temporally fluctuating environmental conditions without plasticity (equation (B49)) is shown by the dashed line. For comparison, purple crosses denote the expected failure of adaptation in static environments in the absence of plasticity. Refer to the caption of figure C6 for an explanation of what the remaining lines represent.

**Figure C14:**
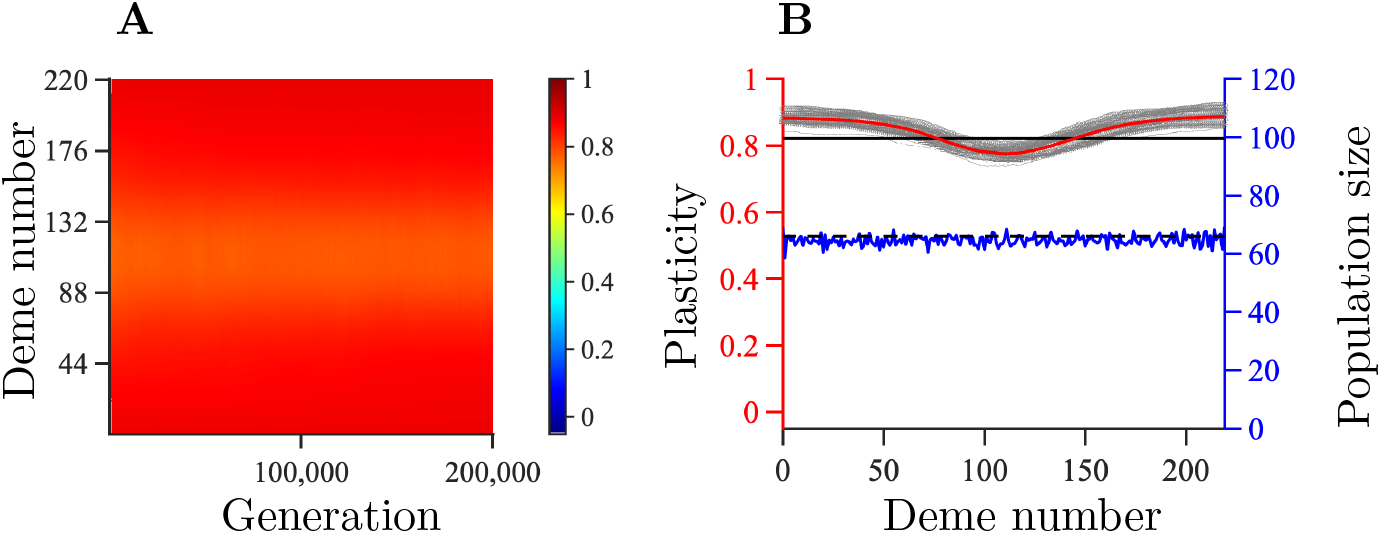
Evolution of plasticity in a habitat with an optimal phenotype that changes linearly in space. Temporal and spatial evolution of plasticity averaged over 100 realisations (A). Population size and plasticity 200, 000 generations after the start of range expansion (B). The red line in panel B shows plasticity averaged over 20 realisations (red axis on the left). The grey area indicates the spread of plasticity between different realisations. The solid black line shows the analytically calculated optimal plasticity. The blue line shows the population size averaged over 20 realisations (blue axis on the right). The black dashed line shows the expected population size for the analytically calculated optimal plasticity (it is overlaid by the solid blue line, indicating good agreement between the simulation and the analytical approximation). Remaining parameters: *θ* = 1.2(*i* − 110.5), *K* = 100, *r*_*m*_ = 1, *V*_*S*_ = 2, *μ* = 10^−6^, *σ* = 1, *L* = 415, *α* = 0.3162, *β* = 0.0024, *γ* = 0.5, *δ* = 0.5, and *σ*_*θ*_ = 0.

**Figure C15:**
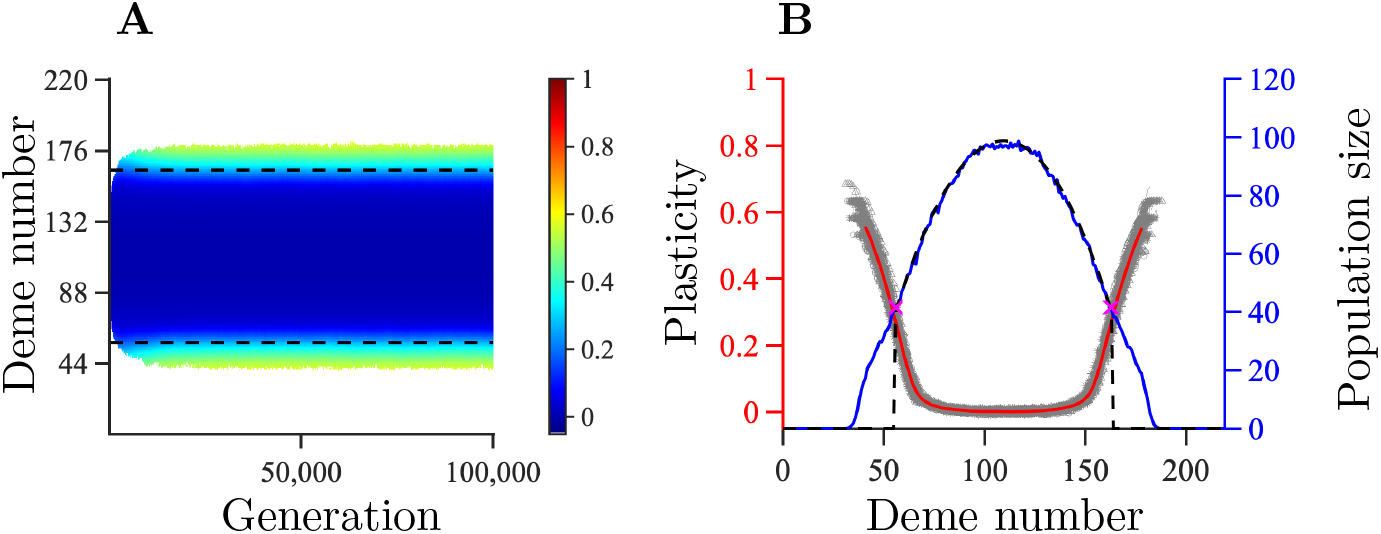
Simulations with the number of loci underlying the plastic component of the trait (*L*_2_) being ten times smaller than the number of loci underlying the non-plastic component of the trait (*L*_1_). Consequently, and in comparison to the results presented in the main text, the allele effect sizes at loci underlying plasticity are here ten times larger than the alleles underlying plasticity in the results presented in the main text. Besides this, all other parameter values correspond to the parameter values shown in figure 2 B. In this case, plasticity that evolves in the range margin is much higher (plasticity of 0.5-0.6) than in the case shown in figure 2 B (plasticity of 0.1). However, because the parameters are within regime *R*_1_, the equilibrium range is finite. Panel A: temporal and spatial evolution of plasticity averaged over 100 realisations. Panel B: population size and plasticity 100, 000 generations after the start of range expansion. Refer to the captions to figures C5 and C6 for explanations of the different lines in panels A and B, respectively. Parameters: *K* = 100, *r*_*m*_ = 1, *V*_*S*_ = 2, *μ* = 10^−6^, *σ* = 1, *L*_1_ = 799, *L*_2_ = 79, *α* = 0.3162, *β* = 0.013, *γ* = 0.5, and *δ* = 0.9.

## References

[1] Sirois-Delisle C, Kerr JT. 2018 Climate change-driven range losses among bumblebee species are poised to accelerate. Scientific Reports 8, 1–10.

[2] Pinsky ML, Selden RL, Kitchel ZJ. 2020 Climate-Driven Shifts in Marine Species Ranges: Scaling from Organisms to Communities. Annual Review of Marine Science 12, 153–179.

[3] Hastings RA, Rutterford LA, Freer JJ, Collins RA, Simpson SD, Genner MJ. 2020 Climate change drives poleward increases and equatorward declines in marine species. Current Biology 30, 1572–1577.

[4] O’Hara CC, Frazier M, Halpern BS. 2021 At-risk marine biodiversity faces extensive, expanding, and intensifying human impacts. Science 372, 84–87.

[5] Yan HF, Kyne PM, Jabado RW, Leeney RH, Davidson LN, Derrick DH, Finucci B, Freckleton RP, Fordham SV, Dulvy NK. 2021 Overfishing and habitat loss drive range contraction of iconic marine fishes to near extinction. Science Advances 7, eabb6026.

[6] Bridle JR, Vines TH. 2007 Limits to evolution at range margins: when and why does adaptation fail?. Trends in Ecology & evolution 22, 140–147.

[7] Sexton JP, McIntyre PJ, Angert AL, Rice KJ. 2009 Evolution and ecology of species range limits. Annual Review of Ecology, Evolution, and Systematics 40, 415–436.

[8] Kirkpatrick M, Barton NH. 1997 Evolution of a species’ range. American Naturalist 150, 1–23.

[9] Chevin LM, Lande R. 2011 Adaptation to marginal habitats by evolution of increased phenotypic plasticity. Journal of evolutionary biology 24, 1462–1476.

[10] Barton NH. 2001 Adaptation at the edge of a species’ range. In Silvertown J, Antonovics J, editors, Integrating ecology and evolution in a spatial context vol. 14 pp. 365–392. Oxford: Blackwell.

[11] Polechová J, Barton NH. 2015 Limits to adaptation along environmental gradients. Proceedings of the National Academy of Sciences 112, 6401–6406.

[12] Sgro CM, Terblanche JS, Hoffmann AA. 2016 What can plasticity contribute to insect responses to climate change?. Annual Review of Entomology 61, 433–451.

[13] Coulson T, Kendall BE, Barthold J, Plard F, Schindler S, Ozgul A, Gaillard JM. 2017 Modeling adaptive and nonadaptive responses of populations to environmental change. The American Naturalist 190, 313–336.

[14] Chevin LM, Hoffmann AA. 2017 Evolution of phenotypic plasticity in extreme environments. Philosophical Transactions of the Royal Society B: Biological Sciences 372, 20160138.

[15] Fox RJ, Donelson JM, Schunter C, Ravasi T, Gaitán-Espitia JD. 2019 Beyond buying time: the role of plasticity in phenotypic adaptation to rapid environmental change. Philosophical Transactions of the Royal Society B: Biological Sciences 374, 20180174.

[16] Scheiner SM, Barfield M, Holt RD. 2020 The genetics of phenotypic plasticity. XVII. Response to climate change. Evolutionary Applications 13, 388–399.

[17] Johansson D, Pereyra RT, Rafajlović M, Johannesson K. 2017 Reciprocal transplants support a plasticity-first scenario during colonisation of a large hyposaline basin by a marine macro alga. BMC Ecology 17, 1–9.

[18] Wang SP, Althoff DM. 2019 Phenotypic plasticity facilitates initial colonization of a novel environment. Evolution 73, 303–316.

[19] Storz JF, Scott GR. 2020 Phenotypic plasticity, genetic assimilation, and genetic compensation in hypoxia adaptation of high-altitude vertebrates. Comparative Biochemistry and Physiology Part A: Molecular & Integrative Physiology p. 110865.

[20] Walter GM, Clark J, Terranova D, Cozzolino S, Cristaudo A, Hiscock SJ, Bridle JR. 2020 Hidden genetic variation in plasticity increases the potential to adapt to novel environments. bioRxiv.

[21] Enbody ED, Pettersson ME, Sprehn CG, Palm S, Wickström H, Andersson L. 2021 Ecological adaptation in European eels is based on phenotypic plasticity. Proceedings of the National Academy of Sciences 118.

[22] Walter GM, Terranova D, Clark J, Cozzolino S, Cristaudo A, Hiscock SJ, Bridle JR. 2021 Plasticity in novel environments induces larger changes in genetic variance than adaptive divergence. bioRxiv.

[23] Tufto J. 2000 The evolution of plasticity and nonplastic spatial and temporal adaptations in the presence of imperfect environmental cues. The American Naturalist 156, 121–130.

[24] Scheiner SM. 2016 Habitat choice and temporal variation alter the balance between adaptation by genetic differentiation, a jack-of-all-trades strategy, and phenotypic plasticity. American Naturalist 187, 633–646.

[25] Schmid M, Dallo R, Guillaume F. 2019 Species’ range dynamics affect the evolution of spatial variation in plasticity under environmental change. American Naturalist 193, 798–813.

[26] Chevin LM, Lande R, Mace GM. 2010 Adaptation, plasticity, and extinction in a changing environment: towards a predictive theory. PLoS Biology 8, e1000357.

[27] Diamond SE, Martin RA. 2021 Buying time: Plasticity and population persistence. In Phenotypic Plasticity & Evolution pp. 185–209. CRC Press.

[28] Ghalambor CK, McKay J, Carroll S, Reznick D. 2007 Adaptive versus non-adaptive phenotypic plasticity and the potential for contemporary adaptation in new environments. Functional Ecology 21, 394–407.

[29] Ghalambor CK, Hoke KL, Ruell EW, Fischer EK, Reznick DN, Hughes KA. 2015 Non-adaptive plasticity potentiates rapid adaptive evolution of gene expression in nature. Nature 525, 372–375.

[30] Gibert P, Debat V, Ghalambor CK. 2019 Phenotypic plasticity, global change, and the speed of adaptive evolution. Current Opinion in Insect Science 35, 34–40.

[31] Leonard AM, Lancaster LT. 2020 Maladaptive plasticity facilitates evolution of thermal tolerance during an experimental range shift. BMC Evolutionary Biology 20, 1–11.

[32] Kellermann V, McEvey SF, Sgr`o CM, Hoffmann AA. 2020 Phenotypic Plasticity for Desiccation Resistance, Climate Change, and Future Species Distributions: Will Plasticity Have Much Impact?. The American Naturalist 196, 306–315.

[33] DeWitt TJ, Sih A, Wilson DS. 1998 Costs and limits of phenotypic plasticity. Trends in Ecology & Evolution 13, 77–81.

[34] Auld JR, Agrawal AA, Relyea RA. 2010 Re-evaluating the costs and limits of adaptive phenotypic plasticity. Proceedings of the Royal Society B: Biological Sciences 277, 503–511.

[35] Scheiner SM, Barfield M, Holt RD. 2017 The genetics of phenotypic plasticity. XV. Genetic assimilation, the Baldwin effect, and evolutionary rescue. Ecology and Evolution 7, 8788–8803.

[36] Scheiner SM. 1998 The genetics of phenotypic plasticity. VII. Evolution in a spatially-structured environment. Journal of Evolutionary Biology 11, 303–320.

[37] Wright S. 1943 Isolation by distance. Genetics 28, 114.

[38] Lande R. 2009 Adaptation to an extraordinary environment by evolution of phenotypic plasticity and genetic assimilation. Journal of Evolutionary Biology 22, 1435–1446.

[39] Eriksson A, Elías-Wolff F, Mehlig B. 2013 Metapopulation dynamics on the brink of extinction. Theoretical Population Biology 83, 101–122.

[40] Eriksson M, Rafajlović M. 2021 The effect of genetic architecture and selfing on the capacity of a population to expand its range. American Naturalist 197, 526–542.

[41] Bridle JR, Kawata M, Butlin RK. 2019 Local adaptation stops where ecological gradients steepen or are interrupted. Evolutionary Applications 12, 1449–1462.

[42] Bulmer MG. 1980 The mathematical theory of quantitative genetics. Clarendon Press.

[43] Bürger R, Lynch M. 1995 Evolution and extinction in a changing environment: a quantitative-genetic analysis. Evolution 49, 151–163.

[44] Chevin LM, Cotto O, Ashander J. 2017 Stochastic evolutionary demography under a fluctuating optimum phenotype. The American Naturalist 190, 786–802.

[45] King JG, Hadfield JD. 2019 The evolution of phenotypic plasticity when environments fluctuate in time and space. Evolution Letters 3, 15–27.

[46] Van Buskirk J. 2017 Spatially heterogeneous selection in nature favors phenotypic plasticity in anuran larvae. Evolution 71, 1670–1685.

[47] Carbonell J, Bilton D, Calosi P, Millán A, Stewart A, Velasco J. 2017 Metabolic and reproductive plasticity of core and marginal populations of the eurythermic saline water bug Sigara selecta (Hemiptera: Corixidae) in a climate change context. Journal of Insect Physiology 98, 59–66.

[48] Tufto J. 2015 Genetic evolution, plasticity, and bet-hedging as adaptive responses to temporally autocorrelated fluctuating selection: A quantitative genetic model. Evolution 69, 2034–2049.

[49] Arnold PA, Nicotra AB, Kruuk LEB. 2019 Sparse evidence for selection on phenotypic plasticity in response to temperature. Philosophical Transactions of the Royal Society B: Biological Sciences 374, 20180185.

[50] Pulliam HR. 1988 Sources, sinks, and population regulation. The American Naturalist 132, 652–661.

[51] Via S, Lande R. 1985 Genotype-environment interaction and the evolution of phenotypic plasticity. Evolution 39, 505–522.

[52] Van Buskirk J, Steiner UK. 2009 The fitness costs of developmental canalization and plasticity. Journal of Evolutionary Biology 22, 852–860.

[53] Murren CJ, Auld JR, Callahan H, Ghalambor CK, Handelsman CA, Heskel MA, Kingsolver J, Maclean HJ, Masel J, Maughan H et al. 2015 Constraints on the evolution of phenotypic plasticity: limits and costs of phenotype and plasticity. Heredity 115, 293–301.

[54] Gavrilets S, Scheiner SM. 1993 The genetics of phenotypic plasticity. V. Evolution of reaction norm shape. Journal of Evolutionary Biology 6, 31–48.

[55] De Jong G. 1999 Unpredictable selection in a structured population leads to local genetic differentiation in evolved reaction norms. Journal of Evolutionary Biology 12, 839–851.

[56] Scheiner SM. 2018 The genetics of phenotypic plasticity. XVI. Interactions among traits and the flow of information. Evolution 72, 2292–2307.

[57] Pigliucci M, Diiorio P, Schlichting CD. 1997 Phenotypic plasticity of growth trajectories in two species of Lobelia in response to nutrient availability. Journal of Ecology pp. 265–276.

[58] Lande R. 2014 Evolution of phenotypic plasticity and environmental tolerance of a labile quantitative character in a fluctuating environment. Journal of Evolutionary Biology 27, 866–875.

[59] Leung C, Rescan M, Grulois D, Chevin LM. 2020 Reduced phenotypic plasticity evolves in less predictable environments. Ecology Letters 23, 1664–1672.

[60] Gabriel W. 2005 How stress selects for reversible phenotypic plasticity. Journal of Evolutionary Biology 18, 873–883.

[61] Lande R. 2015 Evolution of phenotypic plasticity in colonizing species. Molecular Ecology 24, 2038–2045.

[62] Lenormand T, Otto SP. 2000 The evolution of recombination in a heterogeneous environment. Genetics 156, 423–438.

[63] Peck JR, Yearsley JM, Waxman D. 1998 Explaining the geographic distributions of sexual and asexual populations. Nature 391, 889–892.

[64] Fouqueau L, Roze D. 2021 The evolution of sex along an environmental gradient. Evolution.

[65] Leimar O, Dall SR, McNamara JM, Kuijper B, Hammerstein P. 2019 Ecological genetic conflict: genetic architecture can shift the balance between local adaptation and plasticity. American Naturalist 193, 70–80.

[66] Sippel S, Meinshausen N, Fischer EM, Székely E, Knutti R. 2020 Climate change now detectable from any single day of weather at global scale. Nature Climate Change 10, 35–41.

[67] Valenzuela N, Literman R, Neuwald JL, Mizoguchi B, Iverson JB, Riley JL, Litzgus JD. 2019 Extreme thermal fluctuations from climate change unexpectedly accelerate demographic collapse of vertebrates with temperature-dependent sex determination. Scientific Reports 9, 1–11.

[68] Van Oppen MJ, Oliver JK, Putnam HM, Gates RD. 2015 Building coral reef resilience through assisted evolution. Proceedings of the National Academy of Sciences 112, 2307–2313.

[69] Van Oppen MJ, Gates RD, Blackall LL, Cantin N, Chakravarti LJ, Chan WY, Cormick C, Crean A, Damjanovic K, Epstein H et al. 2017 Shifting paradigms in restoration of the world’s coral reefs. Global Change Biology 23, 3437–3448.

[70] Gibbin EM, N’Siala GM, Chakravarti LJ, Jarrold MD, Calosi P. 2017 The evolution of phenotypic plasticity under global change. Scientific Reports 7, 1–8.

[71] Pazzaglia J, Reusch TB, Terlizzi A, Marín-Guirao L, Procaccini G. 2021 Phenotypic plasticity under rapid global changes: The intrinsic force for future seagrasses survival. Evolutionary Applications.

[72] Macić V, Albano PG, Almpanidou V, Claudet J, Corrales X, Essl F, Evagelopoulos A, Giovos I, Jimenez C, Kark S et al. 2018 Biological invasions in conservation planning: a global systematic review. Frontiers in Marine Science 5, 178.

[73] Geng YP, Pan XY, Xu CY, Zhang WJ, Li B, Chen JK, Lu BR, Song ZP. 2007 Phenotypic plasticity rather than locally adapted ecotypes allows the invasive alligator weed to colonize a wide range of habitats. Biological Invasions 9, 245–256.

[74] Davidson AM, Jennions M, Nicotra AB. 2011 Do invasive species show higher phenotypic plasticity than native species and, if so, is it adaptive? A meta-analysis. Ecology Letters 14, 419–431.

[75] Ren GQ, Yang HY, Li J, Prabakaran K, Dai ZC, Wang XP, Jiang K, Zou CB, Du DL. 2020 The effect of nitrogen and temperature changes on Solidago canadensis phenotypic plasticity and fitness. Plant Species Biology 35, 283–299.

[76] Xue Q, Ma CS. 2020 Aged virgin adults respond to extreme heat events with phenotypic plasticity in an invasive species, Drosophila suzukii. Journal of Insect Physiology 121, 104016.

[77] Chown SL, Slabber S, McGeoch MA, Janion C, Leinaas HP. 2007 Phenotypic plasticity mediates climate change responses among invasive and indigenous arthropods. Proceedings of the Royal Society B: Biological Sciences 274, 2531–2537.

[78] Zenni RD, Lamy JB, Lamarque LJ, Porté AJ. 2014 Adaptive evolution and phenotypic plasticity during naturalization and spread of invasive species: implications for tree invasion biology. Biological Invasions 16, 635–644.

[79] Falaschi M, Melotto A, Manenti R, Ficetola GF. 2020 Invasive species and amphibian conservation. Herpetologica 76, 216–227.

[80] Rathee S, Ahmad M, Sharma P, Singh HP, Batish DR, Kaur S, Kaur A, Yadav SS, Kohli RK. 2021 Biomass allocation and phenotypic plasticity are key elements of successful invasion of Parthenium hysterophorus at high elevation. Environmental and Experimental Botany 184, 104392.

[81] Pease CM, Lande R, Bull J. 1989 A model of population growth, dispersal and evolution in a changing environment. Ecology 70, 1657–1664.

[82] Barton N, Turelli M. 1991 Natural and sexual selection on many loci.. Genetics 127, 229–255.

[83] Lande R. 2007 Expected relative fitness and the adaptive topography of fluctuating selection. Evolution 61, 1835–1846.

[84] Rudin W. 1964 Principles of mathematical analysis vol. 3. McGraw-Hill New York.

[85] Slatkin M. 1973 Gene flow and selection in a cline. Genetics 75, 733–756.

